# Evolutionary determinants of non-seasonal breeding in wild chacma baboons

**DOI:** 10.1101/2021.03.18.435953

**Authors:** Jules Dezeure, Alice Baniel, Lugdiwine Burtschell, Alecia J. Carter, Bernard Godelle, Guy Cowlishaw, Elise Huchard

**Author notes:** Corresponding author: Jules Dezeure. Equal contributions (Guy Cowlishaw, Elise Huchard).

## Abstract

Animal reproductive phenology varies from strongly seasonal to non-seasonal, sometimes among closely related or sympatric species. While the extent of reproductive seasonality is often attributed to environmental seasonality, this fails to explain many cases of non-seasonal breeding in seasonal environments. We investigated the evolutionary determinants of non-seasonal breeding in a wild primate, the chacma baboon (*Papio ursinus*), living in a seasonal environment with high climatic unpredictability. We tested three hypotheses proposing that non-seasonal breeding has evolved in response to (1) climatic unpredictability, (2) reproductive competition between females favouring birth asynchrony, and (3) individual, rank-dependent variations in optimal reproductive timing. We found strong support for an effect of reproductive asynchrony modulated by rank: (i) birth synchrony is costly to subordinate females, lengthening their interbirth intervals, and (ii) females delay their reproductive timings (fertility periods and conceptions) according to other females in the group to stagger conceptions. These results indicate that reproductive competition generates reproductive asynchrony, weakening the intensity of reproductive seasonality at the population level. This study emphasizes the importance of sociality in mediating the evolution of reproductive phenology in gregarious organisms, a result of broad significance for understanding key demographic parameters driving population responses to increasing climatic fluctuations.

## INTRODUCTION

Reproduction is energetically costly, and many species therefore adjust the timing of their reproduction, referred to as reproductive phenology, to seasonal fluctuations in food availability (Boyce 1979). Reproductive seasonality, the temporal clustering of reproductive events in one period of the annual cycle (Lindburg 1987), is widespread taxonomically and geographically (Brockman and van Schaik 2005; Bronson 2009), and usually characterized by the timing and width of the birth peak. Although non-seasonal breeders reproduce throughout the annual cycle (unlike seasonal breeders, who experience a period without any births), they may still exhibit a seasonal peak in the annual distribution of births (Janson and Verdolin 2005).

Ultimate explanations for seasonal reproduction have largely assumed that variation in the intensity of reproductive seasonality reflects variation in the intensity of environmental seasonality (Rutberg 1987; Conover 1992; Di Bitetti and Janson 2000). However, numerous sympatric species exhibit a range of reproductive schedules despite sharing the same climate. For example, in the Serengeti National Park (Tanzania), the korrigum *Damascilus korrigum* is a highly seasonal breeder while the phylogenetically related hartebeest *Alcelaphus buselaphus* is not, and seasonal breeders vary widely in the timing and length of their breeding season (Sinclair et al. 2000). More generally, numerous species living in highly seasonal environments breed year round (Burthe et al. 2011; Swedell 2011; Campos et al. 2017). Overall, the intensity of environmental seasonality is not always a reliable predictor of a species’ reproductive seasonality.

Several additional predictors of seasonal breeding could be considered. First, if the height and/or the timing of the annual food peak vary between years, individuals may benefit from maintaining a flexible reproductive schedule (Colwell 1974; van Schaik and van Noordwijk 1985; Loe et al. 2005). For example, across 38 ungulate species, the intensity of seasonal breeding decreases with increasing seasonal unpredictability, i.e., inter-annual variation in the timing and strength of environmental seasonality (English et al. 2012). In environments where within-year (seasonal) variations are negligible compared to between-year (non-seasonal) variations, individuals adjusting their reproductive events with relative flexibility might be favoured to exploit the unpredictable food peak opportunistically (van Schaik and van Noordwijk 1985); this flexibility may cause the absence of reproductive seasonality. In the context of the income-capital breeder continuum, a common framework for the study of alternative strategies to finance offspring production, capital breeders (which are able to store energy for later use) are often better adapted to such between-year variable environments than income breeders (which rely on current energy available to breed) (Drent and Daan 1980; Brockman and van Schaik 2005; Stephens et al. 2014), and as such, capital breeding species may lower the intensity of their reproductive seasonality in response to high between-year environmental variation. Yet, few studies have asked whether seasonal predictability could represent an evolutionary driver of reproductive seasonality.

Second, social factors might further affect reproductive seasonality by modulating reproductive synchrony, the ‘phenomenon caused by biological interactions to produce a tighter clustering of reproductive events than environmental seasonality alone’ (Ims 1990; page 135). Synchronizing births in order to satiate predators is a common anti-predator adaptation producing extreme reproductive synchrony for numerous species, including some ungulates (Rutberg 1987; Ims 1990; Sinclair et al. 2000; Canu et al. 2015) and squirrel monkeys (Boinski 1987). Sociality could also lead to the reverse pattern where reproductive events are staggered to decrease reproductive competition over access to mates, paternal care or food (Wiebe et al. 1995). For instance, oestrus asynchrony has been reported in both seasonal breeding ring-tailed lemurs (*Lemur catta)* (Pereira 1990) and non-seasonal breeding chimpanzees (*Pan troglodytes)* (Matsumoto-Oda et al. 2007), apparently allowing females greater mate choice by decreasing female mating competition.

Third, individual variation in reproductive seasonality might also occur, leading to non-seasonal breeding across the population as a whole. This is especially true in hierarchical societies, where dominant females often have privileged access to resources and may subsequently exhibit earlier age at first reproduction, shorter interbirth intervals, higher offspring survival and increased longevity (Clutton-Brock and Huchard 2013; Stockley and Bro-Jørgensen 2011). The consequences of rank-related variation in foraging success and life history traits on reproductive seasonality have not been studied, but may mediate both the effects of environmental variation and group reproductive synchrony described above. If females compete mainly over food, we should detect rank-related variation in sensitivity to environmental fluctuations. If females compete mainly over mating opportunities, we should detect rank-related variation in sensitivity to group synchrony.

Here, we propose three non-exclusive hypotheses that could explain the evolution of non-seasonal reproduction. (H1) The ‘non-seasonal environment hypothesis’ proposes that the absence of reproductive seasonality stems from a population-level factor, namely non-seasonal (i.e. between-year) environmental fluctuations. (H2) The ‘group asynchrony hypothesis’ proposes that the absence of reproductive seasonality results from a group-level factor, where females within a group stagger their reproductive events to minimize reproductive synchrony in response to reproductive competition. (H3) The ‘social rank hypothesis’ proposes that an individual-level factor, namely social rank, leads to the absence of reproductive seasonality at the population level because there are dominance-related differences in how females are affected by seasonal and non-seasonal environmental variation (H3a), and/or reproductive synchrony (H3b).

Our study model is a wild social primate, the chacma baboon (*Papio ursinus*). Focusing on a long-lived mammal in the tropics will bring a fresh perspective on the breeding seasonality literature, which is biased towards short-lived passerines in northern temperate climates (Verhulst and Nilsson 2008; Bronson 2009; Varpe et al. 2009). In addition, the selective pressures affecting breeding seasonality in tropical latitudes, where the environment is characterized by more unpredictable rainfall (Feng et al. 2013), have been less studied. Extensive variations in patterns of reproductive synchrony occur across primate populations (Ostner et al. 2008; Gogarten and Koenig 2013), and may reflect female reproductive competition (Beehner and Lu 2013). Finally, social primates such as baboons, which live in large multimale-multifemale groups where the female dominance hierarchy is linear and affects foraging success and reproductive performance (Pusey 2012), may provide a valuable model for understanding rank differences in sensitivity to environmental and social factors likely to lead to individual variations in seasonal breeding strategies. In our study population, a recent study revealed fitness variations associated with seasonal birth timing (Dezeure et al. 2021*a*), and hence raises the question of the nature of the benefits of non-seasonal breeding.

We tested these three hypotheses according to their predicted effects on (1) females’ fitness, assayed by: female interbirth interval (IBIs) and offspring survival until weaning; and on reproductive timings, assayed by the monthly probabilities of cycle resumption and conception. Under the non-seasonal environment hypothesis (H1), we predicted that female reproduction should be responsive to current environmental conditions, such that a scarcity of food (after controlling for patterns of seasonal variation) should lead to longer IBIs, higher infant mortality, and lower probabilities of cycling resumption and conception. Under the reproductive synchrony hypothesis (H2), female reproduction should be responsive to the degree of reproductive synchrony in the group, such that higher group synchrony should lead to longer IBIs, higher infant mortality, and lower probabilities of cycling resumption and conception. Under the social rank hypothesis, lower-ranking females should be more restricted in their access to food resources (H3a), or experience greater costs of reproductive synchrony (H3b), such that the respective negative effects of food scarcity and group synchrony on IBI, infant mortality, cycle resumption, and conception would be greater for subordinate females.

## METHODS

### a) Field site and population

Data were collected between 2005 and 2019 from three habituated groups of wild chacma baboons (J and L since 2005, and M, a fission group from J, since 2016) living on the edge of the Namib Desert at Tsaobis Nature Park (22°23S, 15°44’50E) in central Namibia. The Tsaobis environment is characterized by steep rocky hills descending towards alluvial plains, and crossed by the ephemeral Swakop riverbed (Cowlishaw and Davies 1997). It is a strongly seasonal environment: the desert vegetation responds quickly to the austral summer rains, which usually fall between December and April, and then dies back during the dry winter months. The annual rainfall is sparse and highly variable between years (see Figure S1, supplementary material). Each year, a field season of variable length (mean = 137 days, range: 57-240 days) was conducted, mainly during the dry winter season (between May and October). The groups were followed on foot from dawn to dusk on a daily basis, allowing us to collect demographic and behavioural data, as well as GPS locations.

### b) Individual data

A female was considered adult when she reached menarche. The reproductive state of each adult female was monitored on a daily basis. A female could be assigned as: (i) pregnant, with pregnancy being determined *post hoc* following infant birth, and encompassing the six months between the conceptive cycle and the birth; (ii) lactating, for the period until the female resumed cycling after an infant birth; (iii) cycling, including both swollen females in oestrus (i.e., sexually receptive with a perineal swelling) and non-swollen females (i.e. at other stages of their cycle). Conceptive cycles were established based on the beginning of a pregnancy, noticeable with both the paracallosal skin turning red and the absence of other cycles in the following months, and were usually confirmed by a birth. The first post-partum cycle (i.e. cycle resumption) is the first cycle following an infant’s birth, when the female resumes cycling after lactation. The exact date of the cycle resumption corresponds to the first day of oestrus of the first post-partum cycle, i.e. the first day when a sexual swelling is recorded. The dates of these reproductive events were known with accuracy when they were recorded by field observers, and were estimated in their absence using the methods detailed below.

For females born after 2005, their dates of births were either witnessed in the field or estimated with relative precision (see below). For females born before 2005, age was estimated through dentition, using both tooth eruption schedules and the eruption of the molars (Huchard et al. 2009; Kahumbu and Eley 1991).

Females’ parity was determined using long-term life history data and defined as: nulliparous (before the birth of her first infant), primiparous (between the birth of her first and second infant) and multiparous (after the birth of her second infant).

Female social rank was established annually for each group using *ad libitum* and focal observations of agonistic interactions between adult females: supplants, displacements, attacks, chases and threats (Huchard and Cowlishaw 2011). We computed a linear hierarchy using Matman 1.1.4 (Noldus Information Technology, 2013), and then converted to a relative rank to control for group size. Each female was thus assigned one rank per year, ranging from 0 (lowest ranking) to 1 (highest ranking).

Each year, we also computed the number of adult females in each group. A female was considered in the group each year if she was present for more than half of the annual field season.

### c) Individual reproductive data

To test our hypotheses, we considered two measures of female fitness, namely the interbirth interval (‘IBI’) duration and offspring mortality before weaning, and two measures of the timing of female reproduction, namely the monthly probabilities of conception and of cycle resumption.

As baboons are not followed throughout the year at Tsaobis, we had to estimate dates of conceptions, births and cycle resumptions in a number of cases. The dates of those unobserved events—as well as the number of days of uncertainty around those events—were established using a combination of photographs and field observations of infant fur and skin color (ears, eye contours, hands and feet, muzzle, muzzle tip, and ischial callosities), following a protocol detailed in a recent study (Dezeure et al. 2021*b*). In all, we generated a sample of 241 conceptions, with a median uncertainty of 10 days (range: 0-164 days) (Table S1, supplementary material); 215 births, with a median uncertainty of 10 days (range: 0-153 days); and 155 cycle resumptions, with a median uncertainty of 93 days (range: 0-272 days), between 2005 and 2019. The uncertainty of each reproductive event date has been considered in all analyses (see Appendix A, supplementary material).

We defined interbirth intervals (IBIs) as the number of days between two consecutive live births of a mother. We only considered IBIs for which the first infant reached weaning (i.e. survived until 550 days old) (Gesquiere et al. 2017), because females resumed cycling rapidly after their infant’s death (median = 21 days, range = 9-51, n = 9 observed deaths), and their IBIs would have been shortened regardless of environmental or social factors. This age threshold (550 days) was estimated to be the maximum length of post-partum anoestrus in our population (Dezeure et al. 2021*a*), and presumably reflected the upper threshold of weaning age, assuming that females who resumed cycling had weaned their offspring (Lee et al. 1991; Saltzman et al. 2011; Borries et al. 2014). We further removed one IBI where the infant closing the IBI was stillborn (because it might have been due to premature delivery). We computed a total of 120 IBIs from 43 adult females, ranging from 397-1132 days with a mean of 678 days (SD = 128).

For each infant born between 01/01/2005 and 01/08/2018, we investigated whether it died (yes/no) before weaning, i.e. reaching 550 days of age. Death was recorded when a corpse was observed or when the infant had been missing in the group for five consecutive days. Infants born later than August 2018 were not considered as their survival outcome at 550 days was unknown. Four infants that disappeared between two consecutive field seasons (in the absence of observers) were omitted because we could not establish whether the age of death was before or after 550 days. In our final dataset, a total of 39 of 195 infants died before reaching 550 days of age, with mortality occurring at a median age of 74 days (range 1-284 days, n = 17 known dates of death).

### d) Characterization of environmental variation

We first considered four aspects of environmental variation at Tsaobis: the temperature, rainfall, vegetation cover (food availability) and photoperiod (daytime length) (see Appendix B, electronic supplementary material). However, we found that our indicator of food availability (vegetation cover) was always selected as the best environmental variable affecting reproductive success and timings (see Appendices B, C and D, supplementary material, for further details on the selection of this variable). Therefore, we only present in the main text the methods and results associated with food availability.

We used the Normalized Difference Vegetation Index (NDVI) as a measure of vegetation cover and therefore food availability. NDVI is computed using the near-infrared and red light reflected by the surface of an area and measured with satellite sensors; it produces a quantitative index of primary productivity with higher values corresponding to a higher degree of greenness (Didan et al. 2015). It has previously been used as an indicator of habitat quality for the Tsaobis baboons (Baniel et al. 2018*a*; Dezeure et al. 2021*b*) and other baboon populations (Zinner et al. 2001). See Dezeure et al. 2021*a* for more details about our NDVI calculations for each study group.

In subsequent analyses, to test the influence of environmental unpredictability on baboon reproductive seasonality, we disentangled seasonal from non-seasonal variations of NDVI. To achieve this, and for each month of the calendar year, we first computed seasonal variation using the mean monthly NDVI values across all 15 years of study, and labelled this variable ‘NDVI_S’. We then computed non-seasonal variation as the differences between each actual monthly value in a given year and this mean monthly value across years, and labelled this variable ‘NDVI_NS’. For example, for L group in January, NDVI_S=0.140. This refers to the NDVI value observed in January averaged across years throughout the study period (2005-2019). In January 2005, when NDVI=0.158, we calculated NDVI_NS=0.018 (i.e. 0.158 minus 0.140). This positive value of NDVI_NS indicates that the habitat in January 2005 was greener than for an average month of January. More broadly, seasonal variables reflect within-year variation only (predictable variations, consistent between years), while non-seasonal variables reflect between-year variation only (unpredictable, inconsistent between years).

### e) Statistical analyses

We tested our three hypotheses about the causes of non-seasonal breeding (H1, non-seasonal environment hypothesis; H2, group asynchrony hypothesis; H3, individual rank hypothesis) in a series of four models. Models 1 and 2 tested our hypotheses in relation to their effects on female fitness, using two measures: IBI and infant mortality, respectively. Models 3 and 4 tested our hypotheses in relation to their effects on female reproductive timings, again using two measures: the timing of cycle resumption and probability of conception, respectively.

### Models 1 and 2

For Model 1, exploring the length of the IBI following the birth of an infant (in days), we ran a linear mixed model (LMM), while for Model 2, exploring the probability of infant mortality before weaning (i.e. before 550 days), we ran a generalized linear mixed model (GLMM) with a binomial error structure (see Appendix C, supplementary material for more details). The identity of the female/mother was included as a random effect to control for the non-independence of observations of the same female/mother. Both models comprised the following fixed effects. First, to test our hypotheses, we included:

- *Non-seasonal environmental variation (NDVI_NS)*. Under the non-seasonal environment hypothesis (H1), female reproductive performance should be highly sensitive to non-seasonal environmental variations in an environment with important between-year fluctuations of rainfall and food availability. Therefore, we expected females to have longer IBIs and higher offspring mortality before weaning when monthly food availability was lower than average for those months (i.e. accounting for seasonal variation). We estimated the non-seasonal environmental variation using the NDVI_NS variable described above. We averaged NDVI_NS across the whole period spanning the IBI for Model 1, and from conception to 550 days of age (for live infants) or to death (for dead infants) for Model 2.
- *Reproductive synchrony.* Under the asynchrony hypothesis (H2), we expected that when reproductive synchrony increased, IBI would be longer and infant survival would be lower. In both models, we considered as a proxy of reproductive synchrony the number of infants born in the same group, in a given time window of variable length around the birth of the focal infant. We computed the number of infants born *x* months before, *x* months after, and *x* months before and after the birth of the focal infant, where *x* = 1 month, 3 months, or 6 months for a total of 9 variables. We identified the best of these nine variables as the one minimizing the AIC of a model including only this fixed effect (but controlling for all random effects) for each of our two response variables. We then incorporated this best-synchrony variable, which was the number of infants born the past 3 months for IBI, and the 6 following months for infant mortality, into the full model (see Appendix C, electronic supplementary materials for details).
- *Interactions between female rank and both seasonal (see metric below) and non-seasonal (NDVI_NS) environmental variations*. Under the social rank hypothesis (H3a), we expected lower-ranking females to suffer higher fitness costs during environmental harshness.
- *Interaction between female rank and reproductive synchrony*. Under the social rank hypothesis (H3b), we expected lower ranking females to suffer higher fitness costs when reproductive synchrony in the group was higher.

Second, as control variables, we included:

- *Seasonal environmental variation*, to control for those seasonal variations that are known to affect individual fitness, even for this non-seasonal breeder (Dezeure et al. 2021*a*). We captured seasonal environmental variation by using a sine term of the birth date of the focal infant to describe the timing of an infant’s birth in the annual cycle (in radians). We used only one harmonic and changed the phase value φ (to 0, π/6, π/3, π/2, 2*π/3 or 5*π/6), to account for potential phase shifts across the year (see Appendices C and D) (Dezeure et al. 2021*b*). For these models, we could not average NDVI_S to capture seasonal environmental variation as the mean duration between two births (for IBI) and between conceptions and weaning (for mortality) equals two years on average (and thus an average of two years would be constant for every individual). Nonetheless, the sine wave with the best phase was highly correlated with seasonal fluctuations of NDVI (phase = 5* π/6, for J group, Pearson correlation test: R = -0.92, t = -31.5, p < 0.001) so we used only this fixed effect to capture seasonal environmental variation.
- *The number of adult females in the group* during the birth year of the focal infant, which is an indicator of within-group competition. A reduction in group size is often associated with an increase of primate (including baboon) female fertility, in particular by shortening IBI (Altmann and Alberts 2003; Borries et al. 2008).
- *Group identity*, to control for possible differences between social groups.
- *Maternal social rank* during the birth year. Lower-ranking females were expected to have longer IBIs and lower infant survival (Bulger and Hamilton 1987; Packer et al. 1995) (Dezeure at al., in prep.).
- *Maternal parity.* Primiparous females have not yet achieved full body size and may lack the relevant experience to provide optimal offspring care in comparison to multiparous females. Consequently, they could have longer IBIs, and their offspring could face a higher mortality probability (Altmann and Alberts 2005; Gesquiere et al. 2017).
- *Infant sex.* We expected mothers of males to have longer IBIs than mothers of female infants, and possibly lower survival, in this sexually dimorphic species (Bercovitch and Berard 1993; but see Cheney et al. 2004; Gesquiere et al. 2017).

For Model 1, the IBI model, we also included the quadratic effect of female age (years). Following Gesquiere et al. (2017), we expected both younger and older females to have longer IBIs. We kept parity as fixed effect in the IBI model for consistency, and after checking for the absence of collinearity between our fixed effects (parity and female age) in our model (vif < 2).

### Models 3 and 4

In the case of Models 3 and 4, because seasonal reproduction is usually characterized by a mating season (determined by the seasonality of female fertility or sexual receptivity) and/or a birth season (determined by the seasonality of conceptions), we tested our hypotheses in relation to both the monthly timing of cycle resumptions, i.e., the beginning of the mating period for each female (Model 3), and the monthly timing of conceptions (Model 4). We did not analyse births because conceptions and births were highly dependent in our dataset (most conceptions had been estimated based on the dates of birth), and because females should have more flexibility to adjust the timings of conceptions than births.

To calculate the ‘cycle resumption’ response variable (Model 3), we assessed the monthly probability that a female would resume cycling during those time windows in which cycle resumption was possible, i.e., following the post-partum anoestrus period, which lasts between 223 to 550 days in our population (7-18 months). During the 14 years of study and for each female, we considered only those months that were included within this window of possibility (7-18 months after each birth) and coded 0 if she did not commence cycling in a given month, and 1 if she did. Similarly, to calculate the ‘conception’ response variable (Model 4), we considered whether a female was cycling during each month of the 14 years of study, and for each cycling month coded 0 if she did not conceive and 1 if she did. For each model, we ran a GLMM with a binomial error structure, with the identity of the female set as a random effect.

Regarding our predictors, there were several differences in our approach between Models 1 and 2 versus Models 3 and 4, reflecting their different foci on fitness and timing effects respectively. First, we used different proxies of reproductive synchrony in Models 3 and 4 because reproductive timings are likely to be affected by the current and past reproductive states of other females in the group. We used two distinct metrics of reproductive synchrony in these two models, namely the ‘monthly number of conceptions’ and ‘monthly number of cycling females’ in the same group as the focal event. For the first variable, we observed 155 cycle resumptions and 241 conceptions. For the latter variable, we restricted our dataset to those months for which observations were available (to avoid accumulating uncertainties in the date estimates of reproductive state changes), resulting in a dataset comprising n = 61 cycle resumptions and n = 103 conceptions. However, we did not simultaneously include both metrics in the same model, as the numbers of conceptions and cycling females in a given month are correlated (Pearson correlation test: R = 0.23, t = 6.21, p < 0.001). Consequently, we ran Models 3 and 4 twice, using either the number of conceptions (Models 3.1, 4.1) or the number of cycling females (Models 3.2, 4.2) as the measure of reproductive synchrony.

Second, the effects of environmental variations (NDVI_S and NDVI_NS) and group reproductive synchrony (number of cycling females or of conceptions per month) on the timing of cycle resumptions and conceptions may operate over various time periods. We thus used a moving-window approach to consider possible time period effects (van de Pol et al. 2016). For our two environmental variables, we identified the best time window testing periods covering 0 to over 12 months prior to the focal event using an AIC-based selection procedure in a univariate mixed model containing only the fixed effect of interest. We then added the best variable as a fixed effect in our final multivariate model. For the first reproductive synchrony variable, ‘monthly number of conceptions’, we similarly identified the best time window but for different durations, from 0 to 6 months. For the second reproductive synchrony variable, ‘monthly number of cycling females’, we could not explore its effects over the same time periods due to the limitations of our dataset, and we tested only its past and current effect from 0 to 2 month time periods. In all, we thus investigated the effects of the number of conceptions in the same month, the mean number of conceptions over the past 1-6 months, the number of cycling females the same month, and the mean number of cycling females over the past 1-2 months, resulting in 10 different models for each response variable. Details on these procedures are given in Appendix D and Figure S2, supplementary materials.

To test our hypotheses, the fixed effects in both models comprised:

- *Non-seasonal environmental variations (NDVI_NS).* Under the non-seasonal environment hypothesis (H1), we expected the probabilities of cycling resumption and conception to increase when the food availability was higher than average for this particular time of the year.
- *Reproductive synchrony*. Under the asynchrony hypothesis (H2), we expected the probabilities of cycling resumption and conception to increase when the group synchrony decreased.
- *Interactions between female rank and environmental variations (both seasonal and non-seasonal)*. Under the social rank hypothesis (H3a), we expected that higher-ranking females would be less sensitive to environmental variation than lower-ranking ones.
- *Interaction between reproductive synchrony and female rank*. Under the social rank hypothesis (H3b), we expected lower-ranking females only to adjust their reproductive timings depending on group reproductive synchrony.

As control variables we also included:

- *Seasonal environmental variations (NDVI_S)*. Even for non-seasonal breeders, the intensity of the seasonal environment (regardless of its non-seasonal variation) could affect both the probability to resume cycling and to conceive for females (Cheney et al. 2004). In this population, the annual conceptive peak occurs at the end of the rainy season (Dezeure et al. 2021*a*) and we therefore expect seasonal environmental variation to affect conception probabilities.
- *The number of adult females in the group* during the year of the reproductive event. We expected to find a negative effect of this indicator of within-group competition on the probability of conception (Bulger and Hamilton 1987; Beehner et al. 2006; Roberts and Cords 2013), and possibly on cycle resumption timings.
- *Group identity*, to control for possible differences between social groups.
- *Female rank*. Higher-ranking females could exhibit a higher probability of conception, even if this has not been found in previous baboon studies (Wasser et al. 1998; Beehner et al. 2006; Gesquiere et al. 2017).
- *Female parity*. Nulliparous and primiparous females often have lower reproductive performances than multiparous females, in particular a lower probability of conception (Gesquiere et al. 2017).

We did not include interaction terms between rank and environmental variation in those models that used our second measure of reproductive synchrony, namely the number of cycling females, as a fixed effect. This was to avoid overfitting and facilitate model convergence in this restricted dataset.

### Statistical tests

All statistical analysis were conducted in R version 3.5.0 (R Core Team, 2018). To test our hypotheses with our four mixed models, we used the ‘lmer’ (for LMM, i.e. Model 1) or ‘glmer’ (for binomial GLMMs, i.e Models 2, 3 and 4) function of the lme4 package (Bates et al. 2015). The distribution of residuals were checked using the ‘qqPlot’ function of the car package for LMMs (Fox et al. 2019) and using ‘simulateResiduals’ from DHARMa package for binomial GLMMs (Hartig 2020). All quantitative variables were z-transformed to have a mean of zero and a standard deviation of one in order to facilitate model convergence. When the fits obtained were singular, we double-checked the results by running the exact same models with a Bayesian approach, using the ‘bglmer’ function from the blme package (Dorie 2015). To diagnose the presence of multicollinearity, we calculated the variance inflation factor (VIF) for each of the predictors in each full model using the vif function of the R car package (Fox et al. 2019). These VIFs were < 2.5 across all our final models. For each model, in addition to the Wald chi-square tests with associated P-values computed with the ‘Anova’ function of the R package car (Fox et al. 2019), we calculated 95% Wald confidence intervals to assess the strength of the fixed effects. Only those fixed effects whose confidence intervals did not cross zero and whose P-values < 0.05 were treated as having support. Uncertainty in the dates of conceptions, births, and cycle resumptions were taken into account in all models (Appendix A, supplementary material). We also removed from the models presented in the main text the non-significant interactions, in order to interpret the estimates of the simple effects (and when an interaction was significant, we also tested the significance of simple effects, and present the results where necessary).

## RESULTS

We tested our three non-exclusive hypotheses using four models to explain the evolution of non-seasonal breeding, focusing on two fitness parameters (IBI and infant mortality before weaning, Models 1 and 2) and two timings of reproductive events (cycle resumption and conception, Models 3 and 4). See Table 1 for a summary of predictions and results. Below we describe the support for each hypothesis based on the results across the four models.

**Table 1:**
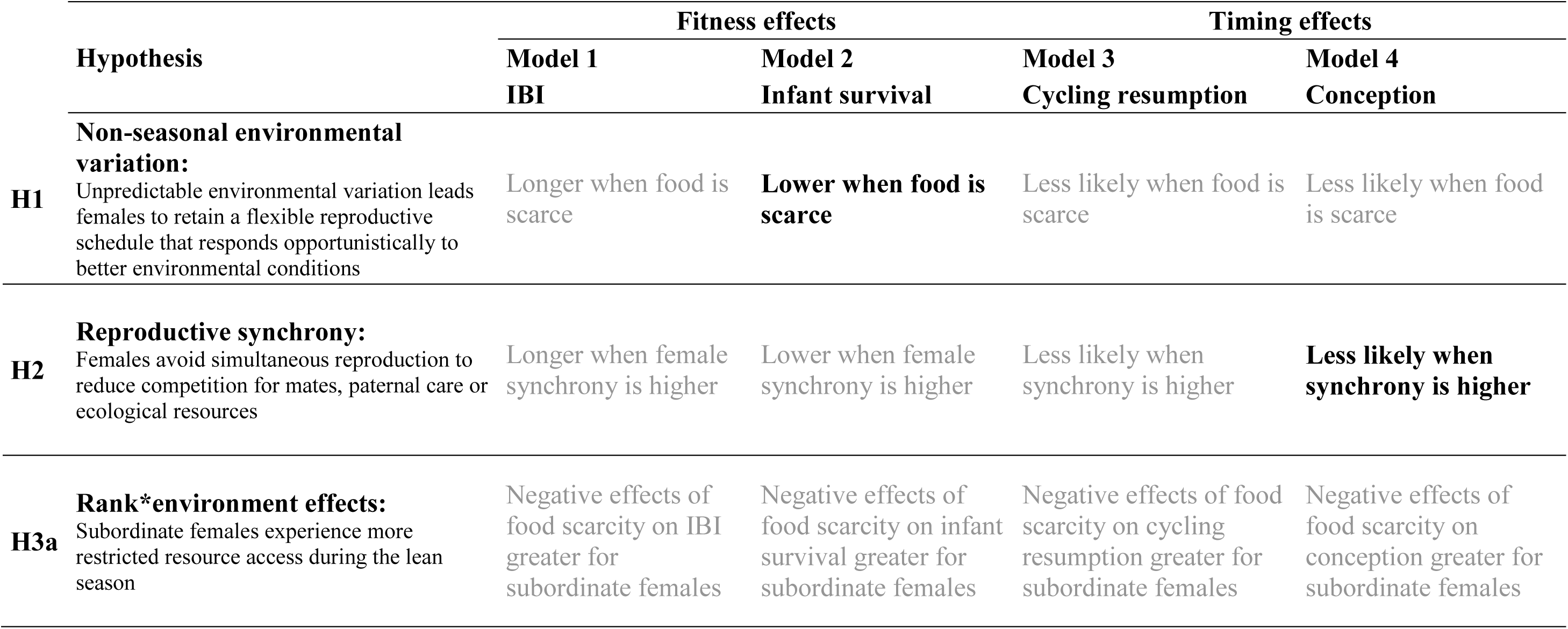
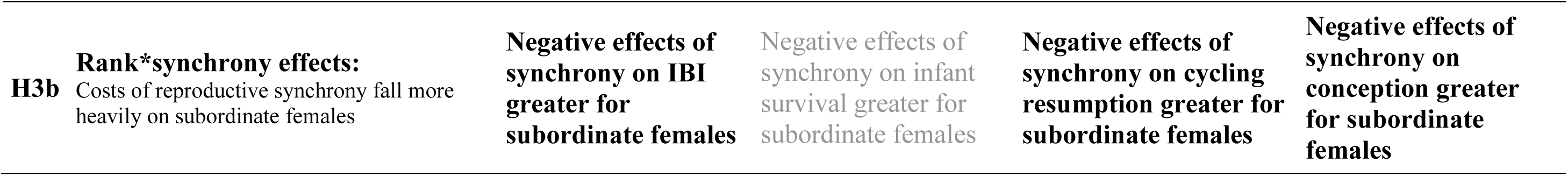
Models and predictions of each fixed effect depending on the hypothesis tested. We indicated with a black and bold writing the predictions supported by our results, and with a grey writing the predictions not supported by our results.

| Hypothesis | Fitness effects |  | Timing effects |  |
| --- | --- | --- | --- | --- |
|  | Model 1<br>IBI | Model 2<br>Infant survival | Model 3<br>Cycling resumption | Model 4<br>Conception |
| **Non-seasonal environmental variation:** |  |  |  |  |
| **H1** Unpredictable environmental variation leads females to retain a flexible reproductive schedule that responds opportunistically to better environmental conditions | Longer when food is scarce | **Lower when food is scarce** | Less likely when food is scarce | Less likely when food is scarce |
| **Reproductive synchrony:** |  |  |  |  |
| **H2** Females avoid simultaneous reproduction to reduce competition for mates, paternal care or ecological resources | Longer when female synchrony is higher | Lower when female synchrony is higher | Less likely when synchrony is higher | **Less likely when synchrony is higher** |
| **Rank\*environment effects:** |  |  |  |  |
| **H3a** Subordinate females experience more restricted resource access during the lean season | Negative effects of food scarcity on IBI greater for subordinate females | Negative effects of food scarcity on infant survival greater for subordinate females | Negative effects of food scarcity on cycling resumption greater for subordinate females | Negative effects of food scarcity on conception greater for subordinate females |

|  |  |  |  |  |  |
| --- | --- | --- | --- | --- | --- |
| **H3b** | **Rank\*synchrony effects:**<br>Costs of reproductive synchrony fall more heavily on subordinate females | **Negative effects of synchrony on IBI greater for subordinate females** | Negative effects of synchrony on infant survival greater for subordinate females | **Negative effects of synchrony on cycling resumption greater for subordinate females** | **Negative effects of synchrony on conception greater for subordinate females** |

### 1/ Minimal support for the non-seasonal environment hypothesis (H1)

Non-seasonal NDVI variation (NDVI_NS) had no effect on our first indicator of reproductive performance: IBI (Table 2). However, survival to weaning was lower for infants who were raised in relatively bad periods in terms of food availability (Table 3, Figure 1), in support of H1. Non-seasonal variation of NDVI did not affect the timing of cycle resumption (Table 4), or probability of conception (Table 5), and thus failed to support H1 in these models. In contrast, we found various effects of seasonal environmental variation, either expressed by the sine term derived from the infant date of birth (for IBI and infant mortality, Tables 2 and 3) or by seasonal NDVI variation (for conception, Table 5 and Figure S3, supplementary material). However, we did not detect any effect of seasonal environment variation on cycle resumption timings (Table 4). In all, we found more effects of seasonal than non-seasonal environmental variation on baboon reproduction.

**Figure 1:**
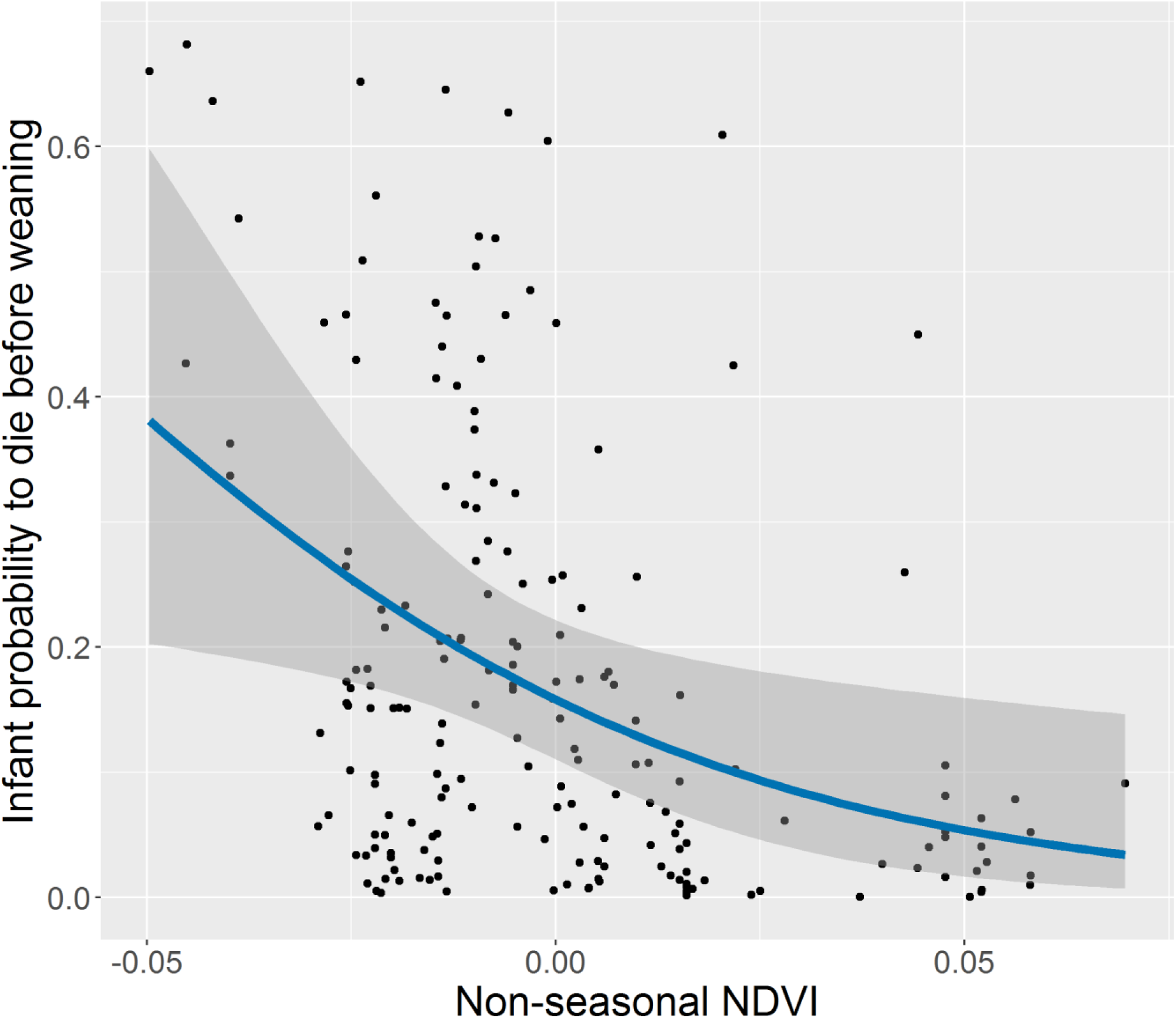
Effects of non-seasonal variations of NDVI on infant mortality before weaning. The black points represent the fitted value of our full model (Table 3) focusing on infant survival before weaning according to the mean value of the non-seasonal NDVI (NDVI_NS) between infant conception and infant death or end of weaning (550 days). The blue curve represents the logistic fit, and the shaded area displays 95% confidence interval around it.

**Table 2:**
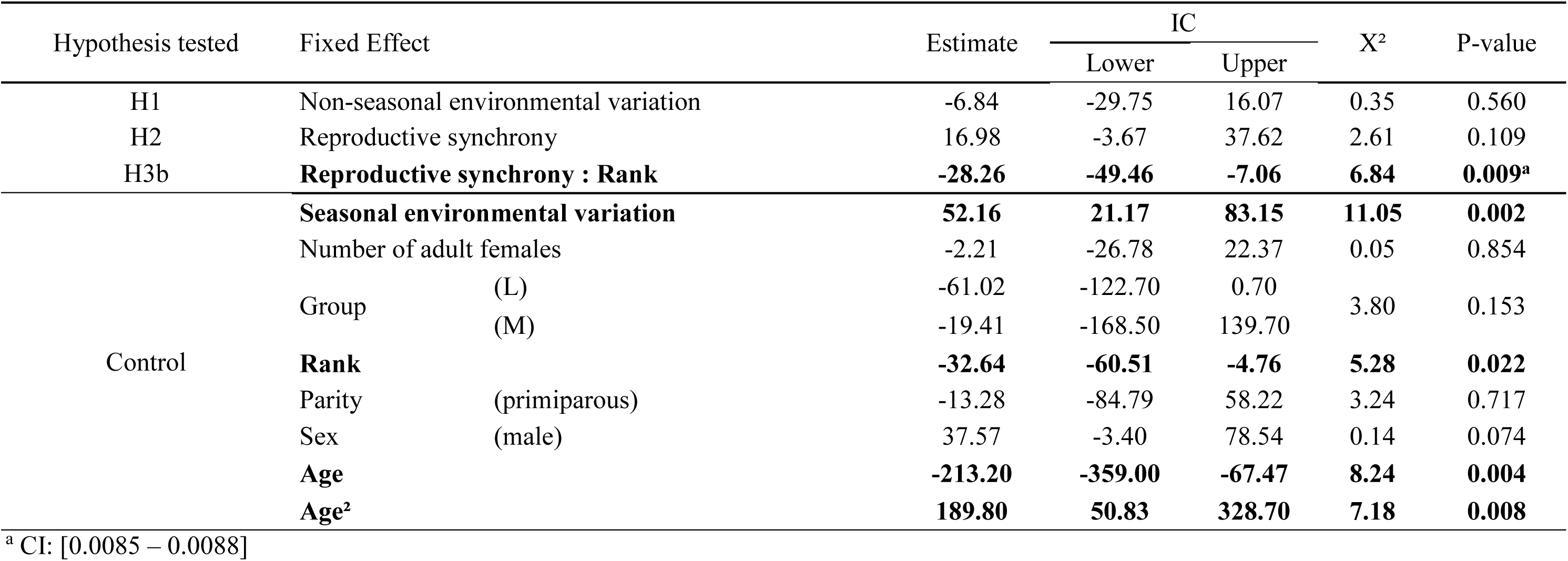
Environmental and social determinants of interbirth interval (IBI) duration (Model 1). We ran 1000 models with simulated birth date, in order to consider the uncertainty of birth dates. The table shows the mean estimates, mean confidence intervals, mean X^2^ statistics and mean p-values of the predictors provided by these 1000 linear mixed models including female identity as random effect, based on 120 observations from 43 females. Significant effects are indicated in bold. For relevant significant effects (i.e. discussed in this study), we also indicated in the footnote the Wilcoxon confidence interval of the p-values. The best fit for seasonal environmental variation is the sine term of the infant date of birth with a phase of π/6. The best fit for non-seasonal environmental variation is the mean ‘NDVI_NS’ between the two births. The best fit for reproductive synchrony is the number of infants born in the group over the past three months. For categorical predictors, the tested category is indicated in parentheses. The interactions between female rank and seasonal environmental variation (X²=0.09, p=0.799), and between female rank and non-seasonal environmental variation (X²=0.47, p=0.496), were not significant and were thus excluded from the full model.

**Table 3:**
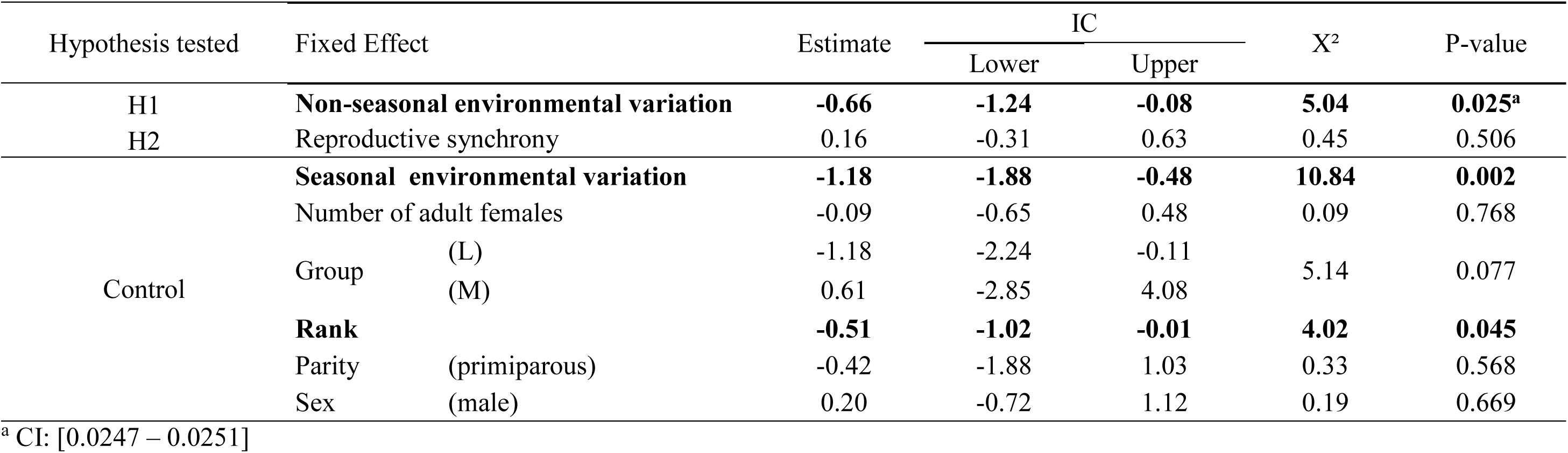
Environmental and social determinants of infant mortality before weaning (0/1: survived until weaning / died before weaning) (Model 2). We ran 1000 models with simulated birth date, in order to consider the uncertainty of birth dates. The table shows the mean estimates, mean confidence intervals, mean X^2^ statistics and mean p-values of the predictors of these 1000 binomial generalized mixed models including female identity as a random effect, based on 19 dead infants out of 195 from 57 females. Significant effects are indicated in bold. For relevant significant effect, we also indicated in the footnote the Wilcoxon confidence interval of the p-values. The best fit for seasonal environmental variation is the sine term of the infant date of birth with a phase of π/2. The best fit for non-seasonal environmental variation is the mean ‘NDVI_NS’ from infant conception to weaning or death. The best fit for reproductive synchrony is the number of infants born in the group the following six months. For categorical predictors, the tested category is indicated in parentheses. The interactions between female rank and seasonal environmental variation (X²=1.89, p=0.186), and between female rank and non-seasonal environmental variation (X²=0.57, p=0.450), were not significant and were thus excluded from the full model.

**Table 4:** Environmental and social determinants on the timing of cycle resumption in a given month, using the number of conceptions as a measure of reproductive synchrony (Model 3.1). We ran 1000 models with simulated cycle resumption date in order to take into account their uncertainty. The table shows the mean estimates, mean confidence intervals, mean X^2^ statistics and mean p-values of the predictors of these 1000 binomial generalized linear mixed model including female identity as a random effect, based on 155 cycle resumptions out of 1768 observations from 56 females. Significant effects are indicated in bold. For relevant significant effect, we also indicated in the footnote the Wilcoxon confidence interval of the p-values. The best fit for seasonal environmental variation is the ‘NDVI_S’ over the past four months. The best fit for non-seasonal environmental variation is the ‘NDVI_NS’ over the past three months. The best fit reproductive synchrony is the mean number of conceptions in the group over the past six months. For categorical predictors, the tested category is indicated in parentheses. The interactions between female rank and seasonal environmental variation (X²=0.33, p=0.555), and between female rank and non-seasonal environmental variation (X²=0.66, p=0.520), were not significant and were thus excluded from the full model.

| Hypothesis tested | Fixed effects | Estimate | IC | | $X^2$ | P-value |
| --- | --- | --- | --- | --- | --- | --- |
|  |  |  | Lower | Upper |  |  |
| H1 | Non-seasonal environmental variation | 0.13 | -0.03 | 0.32 | 2.74 | 0.132 |
| H2 | Reproductive synchrony | -0.20 | -0.37 | 0.01 | 2.16 | 0.192 |
| H3b | **Reproductive synchrony : Rank** | **0.22** | **0.03** | **0.41** | **5.23** | **0.036<sup>a</sup>** |
| Control | Seasonal environmental variation | -0.13 | -0.31 | 0.05 | 2.46 | 0.202 |
|  | Number of adult females | -0.03 | -0.31 | 0.26 | 0.18 | 0.749 |
|  | Group (L) | 0.14 | -0.22 | 0.49 | 1.66 | 0.453 |
|  | Group (M) | -0.61 | -1.95 | 0.73 |  |  |
|  | Rank | 0.04 | -0.13 | 0.21 | 0.93 | 0.795 |
|  | Parity (primiparous) | -0.23 | -0.68 | 0.23 | 0.97 | 0.330 |

<sup>a</sup> CI: [0.02719 – 0.0308]

**Table 5:**
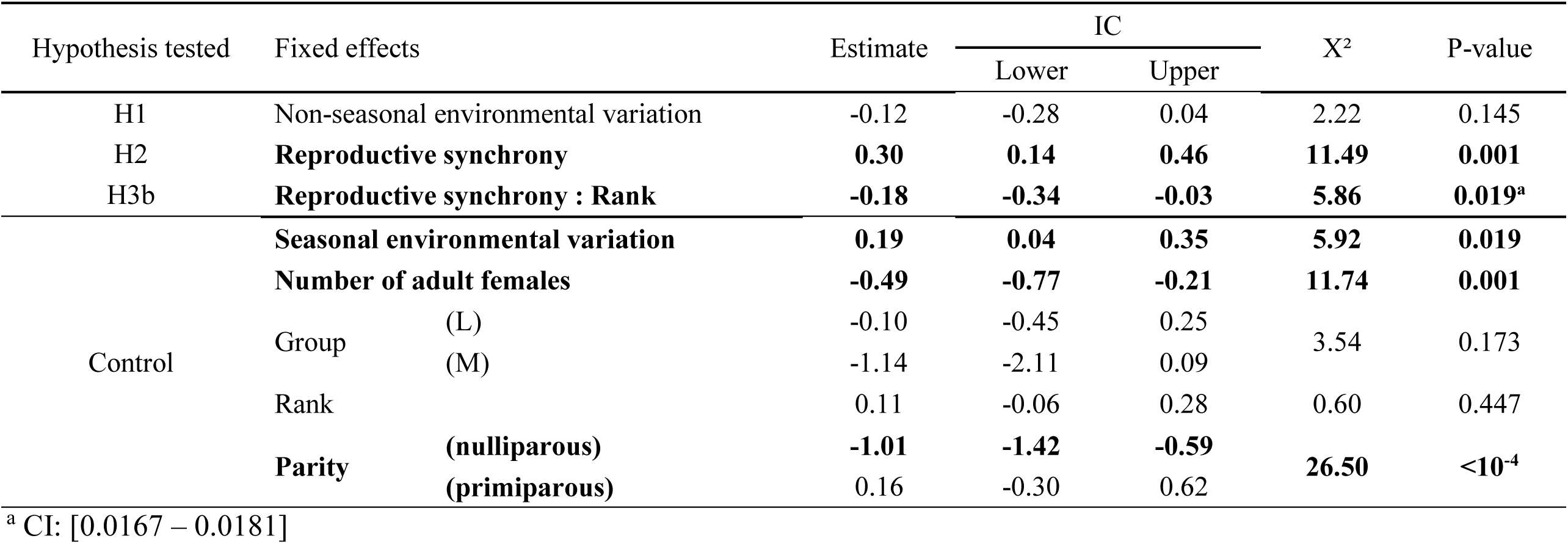
Environmental and social determinants on the probability of conception in a given month, using the number of conceptions as a measure of reproductive synchrony (Model 4.1). We ran 1000 models with simulated conception date in order to take into account their uncertainty. The table show the mean estimates, mean confidence intervals, mean X^2^ statistics and mean p-values of the predictors of these 1000 binomial generalized linear mixed model including female identity as a random effect, based on 241 conceptions out of 1484 observations from 68 females. Significant effects are indicated in bold. For relevant significant effect, we also indicated in the footnote the Wilcoxon confidence interval of the p-values. The best fit for seasonal environmental variation is the ‘NDVI_S’ over the past two months. The best fit for non-seasonal environmental variation is the ‘NDVI_NS’ over the past 12 months. The best fit reproductive synchrony is the mean number of conceptions in the group over the past four months. For categorical predictors, the tested category is indicated in parentheses. The interactions between female rank and seasonal environmental variation (X²=0.07, p=0.842), and between female rank and non-seasonal environmental variation (X²=0.84, p=0.821), were not significant and were thus excluded from the full model.

### 2/ Support for the reproductive asynchrony hypothesis (H2)

We found no effect of reproductive synchrony on IBI (Table 2), infant mortality (Table 3), or the timing of cycle resumption (whether synchrony was measured by the number of cycling females or the number of conceptions, Tables S2 and Table 4, respectively), contrary to H2. However, the influence of reproductive synchrony on the probability of conception showed mixed results depending on the measure of synchrony employed: the number of cycling females had a negative effect, in line with H2 (Figure 2, Table S3), but the number of conceptions had a positive effect, contrary to H2 (Table 5; without the interaction between rank and reproductive synchrony, estimate = 0.28, X^2^ = 11.64, p-value <0.001, CI p-value = [0.00075 – 0.00085]). More precisely, we detected a positive effect of the mean number of monthly conceptions over the past 4 months on the probability to conceive. The same effect was detected over the past 2, 3, 5 and 6 months, but not the same month or previous month (Table S4, supplementary material). This result indicates that female conceptive probability responds more to past than current numbers of conceptions in the group. Thus, we find evidence for a lagged effect of reproductive synchrony, where greater reproductive synchrony, i.e. a higher number of cycling females and higher number of conceptions in the group, suppresses current conceptions in others who are then more likely to conceive once that suppressive effect has been released in later months.

**Figure 2:**
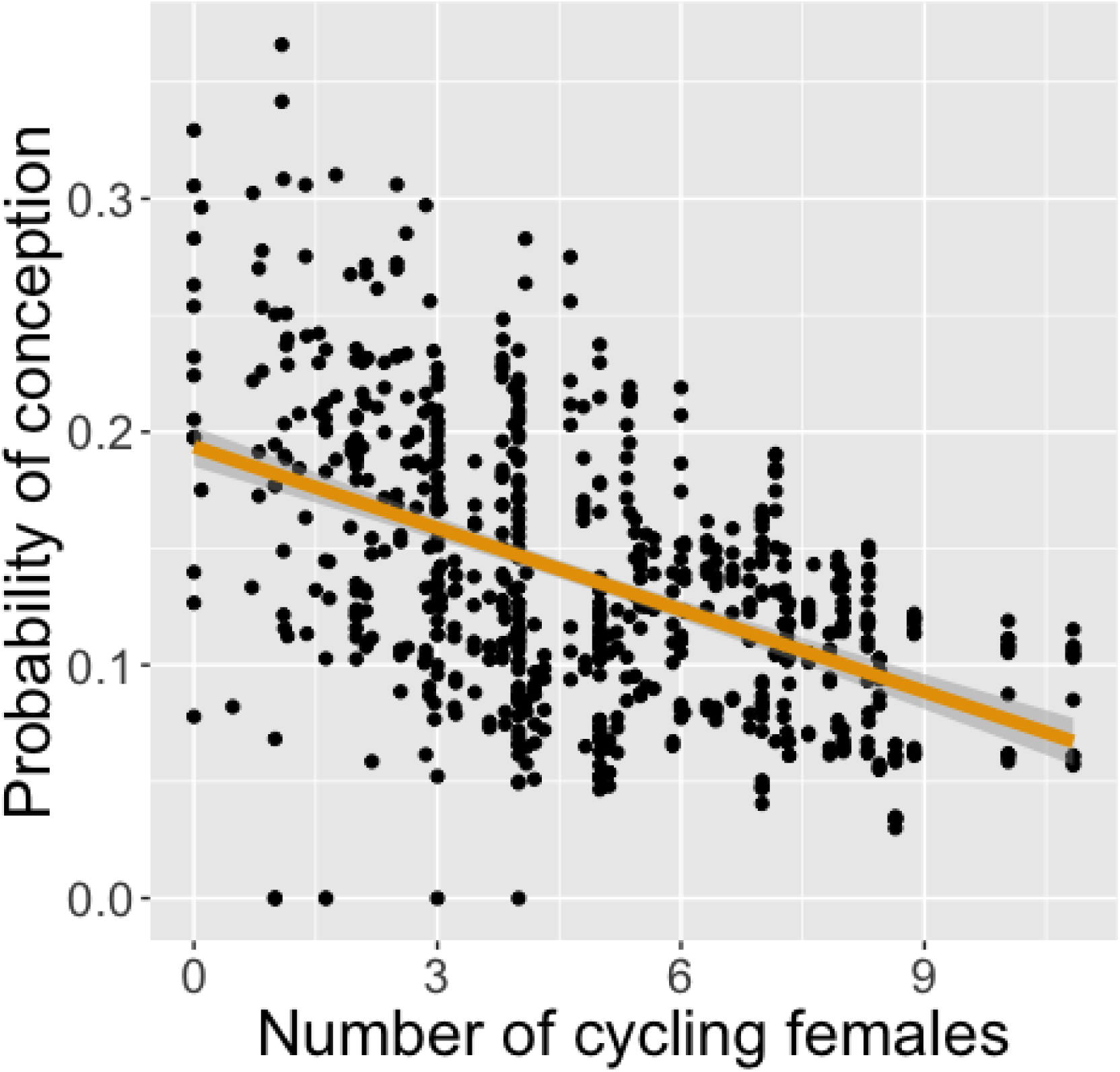
Negative effects of the number of cycling females in the group on the likelihood to conceive in a given month. The black points represent the fitted value of our full model (Model 4.2, Table S3) focusing on the effects of one indicator of group reproductive synchrony, i.e. the number of cycling females in the group, on the conception likelihood. The orange curve represents the linear fit, and the shaded area displays 95% confidence interval around it.

### 3/ Social rank hypothesis: strong support for rank-related variation in response to reproductive synchrony (H3b), but not in response to environmental fluctuations (H3a)

We did not detect any rank-related variation of reproductive seasonality in response to non-seasonal environmental fluctuations (H3a), tested by our interactions between rank and environmental variation for each of our four measures of reproduction: IBI (Table 2), infant survival (Table 3), the timing of cycle resumption (Tables 4), and the probability of conceptions (Table 5). However, we did find support for rank-related variation in reproductive seasonality in response to reproductive synchrony (H3b), tested by interactions between rank and synchrony, in three of our four measures: IBI (Table 2) and the timing of cycle resumption and probability of conception when synchrony was indexed by the number of conceptions (Tables 4 and 5 respectively); such effects were not seen in infant survival (Table 3) or either the timing of cycle resumption or probability of conception when synchrony was indexed by the number of cycling females (Tables S2 and S3 respectively). We will consider each of these three observed effects in turn. Firstly, subordinate females experienced longer IBIs when more infants were born in the group in the three months before they gave birth; this effect was not detectable for high-ranking females (Table 2, Figure 3). Secondly, lower-ranking females were less likely to resume cycling when there had been more conceptions over the past 6 months, whereas higher-ranking females were unaffected (Table 4, Figure 4A). This interaction was not significant for other time windows (number of conceptions occurring in the past 0-5 months before the focal cycle resumption) (Table S5, supplementary material). Thirdly, subordinate females were more likely to conceive when there had been more conceptions in the group over the past four months but this pattern was not seen in dominant females (Table 5, Figure 4B). The same effect was also detected over the past 1, 3 and 5 months, but not 2 and 6 months nor the same month (Table S4, supplementary material). Therefore, subordinate females were more likely to have delayed conceptions, generating breeding asynchrony in the group.

**Figure 3:**
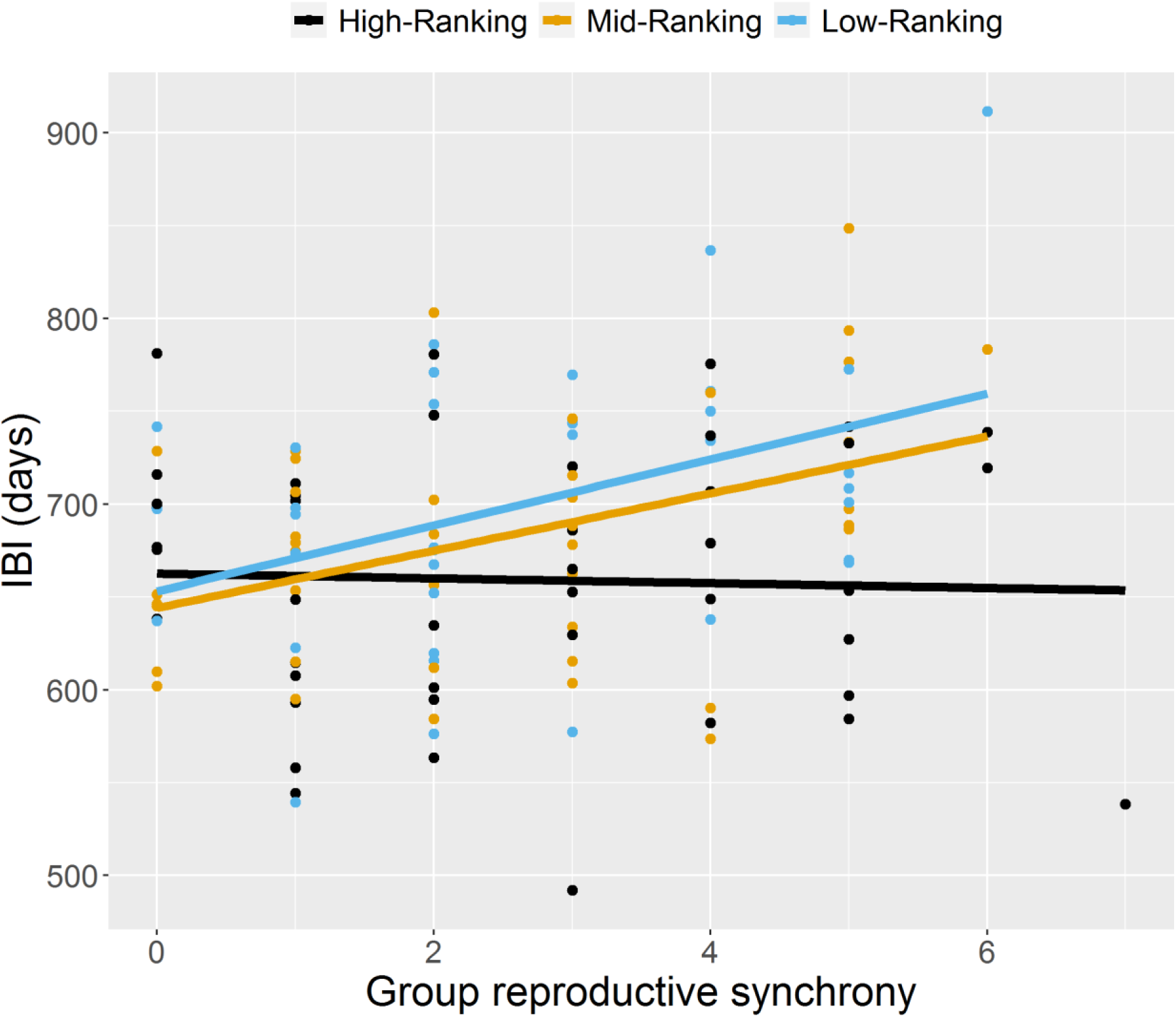
Low and mid-ranking females, but not high-ranking ones, show longer interbirth intervals (IBI) when reproductive synchrony in their group is higher. Each dot represents a fitted value of the full model (Table 2) focusing on IBI according to the number of infants born in the focal female’s group in the three months preceding the birth of her infant. For illustrative purposes, female social rank has been converted into a categorical variable, with females being low-ranking when their rank value is below 0.34 (blue points), high-ranking when it is above 0.67 (black points), and mid-ranking otherwise (orange points). The three curves represent the linear fit for each social rank.

**Figure 4:**
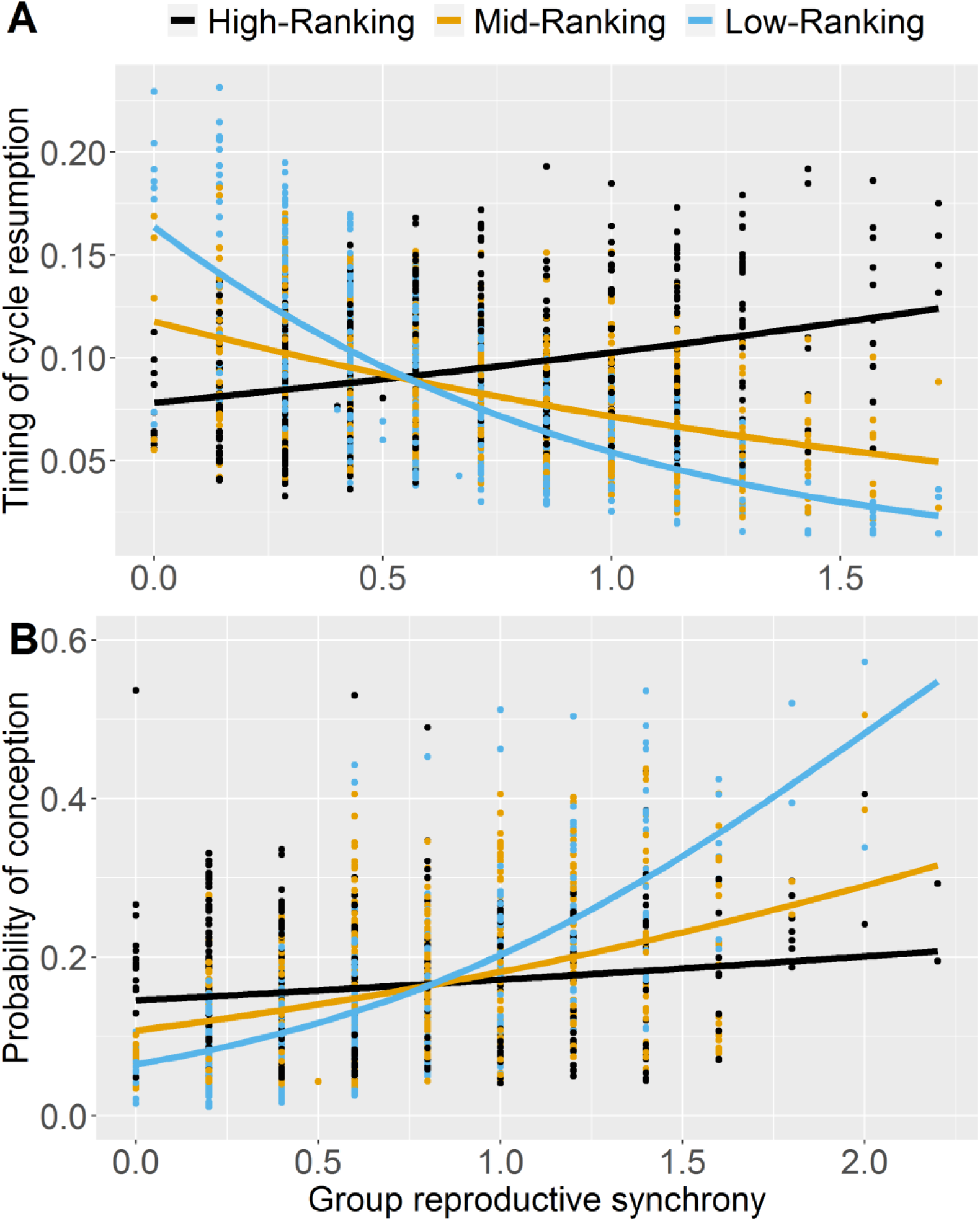
Timings of cycle resumption and monthly probability of conception of lower-ranking females are affected by group reproductive synchrony. The group reproductive synchrony is either the mean number of conceptions over the past 6 months for the timing of cycle resumption (Panel A), or the mean number of conceptions over the past 4 months for the probability of conception (Panel B). For illustrative purposes, female social rank has been converted into a categorical variable, with females being low-ranking when their rank value is below 0.34 (blue points), high-ranking when it is above 0.67 (black points), and mid-ranking otherwise (orange points). Each dot represents a fitted value of the full model focusing on timing of cycle resumption (Model 3.1, Table 4) or probability of conception (Model 4.1, Table 5). The coloured curves show the logistic fit for different categories of social ranks, using the glm method of stat_smooth function of ggplot2 package.

## DISCUSSION

Our study emphasizes the importance of the social environment, and more precisely of group reproductive synchrony, on reproductive phenology in a wild social primate. Females’ reproductive events are staggered, avoiding synchronous breeding, regardless of the season. Below we speculate about the potential drivers of the intrasexual reproductive competition driving breeding asynchrony, with limited access to paternal care as a primary candidate. We further shed light on individual variation in strategies over reproductive phenology dependent on female rank.

### The asynchrony hypothesis to explain non-seasonal breeding (H2 and H3b)

Several results converged to indicate that non-seasonal breeding was an emergent consequence of individual strategies aimed at limiting reproductive competition, leading to breeding asynchrony. **First**, our results showed a fitness cost of birth synchrony, for subordinate females only, who experienced a longer interval to the next conception when more infants were born in the same group in the three months preceding the birth of their own infant. Such an effect is likely to be the consequence of delayed cycle resumption and hence conception experienced by subordinate females when breeding in synchrony with other females of the group. **Second**, the more conceptions that occurred during the past six months in the group, the less likely it was that a lower-ranking female would resume cycling. Such an effect could potentially result in the total loss of a seasonal birth peak at the population level by mechanistically staggering conceptions. In line with this, there was no effect of seasonal or non-seasonal environmental variation on the probability of cycling resumption, indicating that reproductive competition may be the most important factor causing the extension of a mating season in chacma baboons, and thus decreasing the strength of reproductive seasonality. **Third,** our results further showed that the chances of conception decreased when the number of cycling females in the group increased, irrespective of dominance rank. The same pattern was observed with the number of adult females. Both patterns will serve to extend periods of sexual receptivity – and thus lengthen the mating season – across all females. **Fourth**, the likelihood to conceive increased after conceptions had peaked in a female’s group in the previous months, but was unaffected by the number of conceptions occurring during the same month. Subordinate females were more strongly affected by this positive effect. Although a positive effect of others’ conceptions on a female’s conceptive probability was unexpected, it may reflect reproductive asynchrony rather than synchrony, where females, in particular lower-ranking ones, conceive after others have conceived. Because conceptions, unlike cycle resumptions, are influenced by the season, with a moderate peak between April and May, such delays in conceptions likely help to flatten and extend this peak at the group level.

At the proximate level, the reproductive suppression of subordinates by dominants can be explained by female-female competition at different stages of the reproductive cycles. Previous evidence in this population shows that females face increased aggression when they are cycling at the same time as other females (Huchard and Cowlishaw 2011; Baniel et al. 2018*a*). Another recent study suggests that the stress induced by female-female aggression may down-regulate female reproductive physiology, as the harassment of cycling females by other females decreased the likelihood of conception for the victim in a form of reproductive suppression (Baniel et al. 2018*b*). At the ultimate level, females likely adopt competitive tactics to minimise the number of other females with whom they will have to share paternal care in a polygynous mating system, where the alpha male sires nearly 70% of offspring born in the group (Tsaobis: Huchard et al. 2010; other baboons: Alberts 2012). Such an interpretation is strengthened by recent evidence that pregnant and lactating females aggressively target cycling females who mate with the father of their offspring, presumably in an attempt to delay the birth of a paternal half-sibling (Baniel et al. 2018*b*). Paternal care in baboon societies increases offspring survival and growth (Charpentier et al. 2008) through several mechanisms, including protection against infanticide (Huchard et al. 2010; Palombit 2012; Palombit 2003; Palombit et al. 1997), aggression from conspecifics (Lemasson et al. 2008; Nguyen et al. 2009), and increased access to food during weaning (Huchard et al. 2012). Another ultimate explanation may be that females and weanlings compete over present or future access to food. However, the predictions generated by these hypotheses, such as the fact that we should observe more aggression among lactating females than among females in other reproductive states, more aggression among females at times of food scarcity, or higher infant mortality when birth synchrony is high, have all received weak support in our population (Huchard and Cowlishaw 2011; Baniel et al. 2018*a*; this study).

More generally, although fitness costs of synchronous births have been reported in several species, including social primates like yellow baboons (Wasser and Starling 1988) and bonnet macaques (Silk 1989), cooperative breeders such as meerkats and banded mongooses (Clutton-Brock et al. 2001; Nichols et al. 2012), and polygynous birds where females compete over paternal care (Yasukawa et al. 1990, 1993; Slagsvold and Lifjeldt 1994), their consequences for patterns of seasonal reproduction have never been established at the population level. Our study provides the first empirical evidence that reproductive competition at the individual level may generate reproductive asynchrony at the population level.

### Weak support for the non-seasonal environment hypothesis (H1) and no support for a rank-mediated response to seasonal and non-seasonal environmental variations (H3a)

Periods of unusual food scarcity (beyond the seasonal norm) increased offspring mortality before weaning. Food scarcity is known to negatively impact offspring survival before weaning in baboon and other primate populations (Altmann et al. 1977; Cheney et al. 2004; Kleindorfer and Wasser 2004; Gogarten et al. 2012; Campos et al. 2020). Nonetheless, these studies did not disentangle predictable/seasonal from unpredictable/non-seasonal environmental variation, whereas our results show that when infants face harsher periods in early life than usual (i.e. lower than the yearly average), they are less likely to survive. Extreme climatic conditions, including severe droughts, become more frequent with climate change (Easterling et al. 2000; Dai 2013), and our results confirm, in addition to other studies (Wiederholt and Post 2011; Korstjens and Hillyer 2016; Campos et al. 2020), that extreme climatic conditions would negatively affect the demography of wild primates – and probably many other taxa.

We failed to detect any additional evidence in support of the non-seasonal environment hypothesis. Females did not adjust their cycle resumptions or conceptions in response to between-year variations of rainfall or food availability, but only adjusted their conceptions according to seasonal variations in food availability (Tables 4 and 5). Despite important year-to-year variation in cumulative rainfall and thus food availability in Southern Africa, rainfall nearly always occurs during the rainy season (Figure S1; Alberts et al. 2005). In other words, the timing of rains, when they occur (and the associated food peak), remains relatively predictable, which may explain why female reproductive phenology is more responsive to seasonal than non-seasonal environmental variations. In addition, we did not detect any effect of non-seasonal environmental variation on IBI duration, in contrast to seasonal variation (Table 2). This result may reflect the fact that within an IBI, different reproductive stages likely have different sensitivities to past and present environmental variation ; this may be complicated by the fact that IBIs are long at Tsaobis, and so may often integrate both better-than-average and worse-than-average periods, which may cancel each other out. It would possibly limit our ability to detect the impact of year-to-year environmental variation in such long-lived species with our 15-year sample in this study.

Females show similar reproductive responses to variation in food availability irrespective of their social rank. It is possible that subordinate females use more non-monopolisable and/or fallback foods during food scarcity, or negotiate the tolerance of dominant females to access food patches (Sick et al. 2014; Marshall et al. 2015). This ability of subordinates to develop counter-strategies to limit the costs of contest feeding competition has been found in a wide range of animal species (Bugnyar and Kotrschal 2004; Hewitson et al. 2007; Held et al. 2010). Nevertheless, although subordinate females might have been able to minimise competition for food, it appears they were less able to minimise competition over access to mates, given the observed rank-dependent effects of reproductive synchrony on female reproduction. This may reflect the fact that subordinate females can spread out and forage away from dominant females to minimise feeding competition, but cannot do the same for mating competition because dominant females tend to be closely associated with males.

### Other perspectives to explain the evolution of non-seasonal breeding

On top of reproductive asynchrony, several additional ecological or evolutionary mechanisms can help to explain non-seasonal breeding in social primates, and probably beyond. First, a previous study in this population showed the existence of two optimal birth timings in the annual cycle, because the timing that minimizes the mother’s IBI occurs four months before the timing that minimizes offspring mortality (Dezeure et al. 2021*a*). Maternal trade-offs over birth timing may be widespread, and help to explain extended birth seasons between these optima due to individual variation in trade-off resolution. Second, offspring mortality occurs year-round for a variety of reasons including disease, predation, and infanticide; the latter cause can affect up to 30% of offspring in some baboon populations (Palombit 2012). Females typically resume reproduction quickly after losing an infant in social primates (Palombit et al. 2000; Palombit 2003, 2012, 2015) and this may occur even outside the reproductive season. For example, in geladas (*Theropithecus gelada*), females who lose an infant from infanticide following male takeovers can conceive outside the main mating season, resulting in a second annual peak of births, as male takeovers are seasonal (Tinsley Johnson et al. 2018). Such ability to conceive/resume reproduction year-round, which may have evolved in response to high levels of infanticide or more broadly extrinsic infant mortality (infanticide spans from 19 to 28% in our population: Baniel et al. in prep.), may contribute to the absence of seasonal breeding at the population level. Future studies could usefully explore the interplay between ecology, life-history and sociality that may influence variation in the intensity of reproductive seasonality across mammals, a field that has been largely understudied.

### Conclusion

We detected fitness costs associated with birth synchrony, and showed that the reproductive phenology of females, in particular subordinates, is influenced by the phenology of other females in the group in a way that minimizes the costs of synchronous births, which are likely linked to intrasexual competition over access to paternal care. In contrast, female reproductive events were poorly predicted by unpredictable variations in resource availability, such as droughts or peaks of primary productivity, suggesting that females have their reproduction timed according to future, predictable food peaks, rather than relying on the energetic stores accumulated from past unpredictable food availability. This study highlights the importance of female reproductive competition as a main driver of the evolution of reproductive seasonality in social mammals, opening new avenues for future research. It further points to the necessity of understanding the causes and mechanisms of competition at the behavioural level to gain insights on key demographic parameters of a population, including its responses to environmental change.

## ACKNOWLEDGMENTS

This research was carried out with the permission of the Ministry of Environment and Tourism, the Ministry of Land Reform, and the National Commission on Research, Science, and Technology. We further thank the Tsaobis beneficiaries for permission to work at Tsaobis, the Gobabeb Namib Research Institute for affiliation, and Johan Venter and the Snyman and Wittreich families for permission to work on their land. This study was funded by a grant from the Agence Nationale de la Recherche (ANR ERS-17-CE02-0008, 2018-2021) awarded to EH. This paper is a publication of the ZSL Institute of Zoology’s Tsaobis Baboon Project. Contribution ISEM n°XX.

## Authors’ contribution statement

JD, GC and EH conceived the ideas and designed the methodology; JD, AB, AJC, GC, EH collected the data; JD analyzed the data; JD led the writing of the manuscript. All authors contributed critically to the drafts and gave final approval for publication.

## DATA, CODE AND SUPPORTING MATERIAL

Data and codes used in this study are provided in the following GitHub repository: https://github.com/JulesDezeure/Evolutionary-Determinants-of-non-seasonal-breeding-in-baboons.

