## Supplementary Material for "Evolutionary determinants of non-seasonal breeding in wild chacma baboons"

†Equal contributions

**TABLE S3:** ENVIRONMENTAL AND SOCIAL DETERMINANTS ON THE PROBABILITY OF CONCEPTION IN A GIVEN MONTH, USING THE NUMBER OF CONCEPTIONS AS A MEASURE OF REPRODUCTIVE SYNCHRONY (MODEL 4.2)... 21

### APPENDICES

#### Appendix A. Uncertainty of the dates of conceptions, births and cycle resumptions in our models

Dates of conceptions, births and cycle resumptions were estimated in many cases, because the Tsaobis baboons were not followed all year long. Uncertainty in these estimations varied with the time of year, as we generally followed baboons during the cold and dry months, which introduced a systematic bias in our dataset that was important to account for in these analyses. We ran a set of randomizations to evaluate the robustness of the significant effects (see also Dezeure et al. 2021*a*, following the same procedure). For each estimated value of our variables (date of cycle resumptions, conceptions and births), we created an artificial set of 1000 simulated dates covering the full range of potential dates for a given reproductive event. A random date was drawn between the minimal and maximal estimate for each value of a given reproductive parameter, and this procedure was repeated 1000 times. For example, for a birth estimated to occur between the $13^{th}$ of January and the $2^{nd}$ of February, we randomly chose a date between Jan $13^{th}$ and Feb 2nd in one given iteration of our 1000 simulations.

For (i) the costs and benefits of reproductive seasonality on IBI and offspring mortality, the randomizations affected the sine term of the focal infant's date of birth, introduced as a fixed effect in the full model, as these dates of birth could be uncertain. We ran 1000 models, and computed the mean estimates, confidence intervals, $X^2$ tests and p-values associated based on these simulations. For (ii) the influence of social and ecological factors on reproductive seasonality, the randomization affected the response variables (probability to resume cycling and to conceive). We similarly ran 1000 models using each of the 1000 simulated dataset, and extracted the mean estimates, confidence intervals, $X^2$ test and p-values associated. All tables of model results (Table 2-6) presented in main text indicate the mean values of the parameters obtained through these randomization process in order to control for uncertainties in the estimates of reproductive events. We further computed a 95% confidence interval for the relevant p-values of each model, i.e. significant or borderline effects in support of our hypotheses, using a Wilcoxon test. These p-values' confidence intervals are shown in the footnotes of the Tables presented in the main text.

**Appendix B. Characterization of Tsaobis environmental variation**

We considered four environmental aspects to characterize environmental variation at Tsaobis: the photoperiod (daytime length), temperature, rainfall and vegetation cover (food availability). First, we extracted these four environmental factors as follows:

(i) Daily daytime length at Karibib (situated 60km north of Tsaobis) was computed using sunset and sunrise time in this town from a website ('https://dateandtime.info'), then converted into minutes and averaged per month.

(ii) Mean daily daytime land surface temperatures over an 8-day period per pixel of 1km*1km resolution were obtained using MODIS data (product MOD11A2) provided by NASA (Wan et al. 2015). These data were extracted from a rectangular geographic area encompassing the global ranging area of the Tsaobis baboons, computed using GPS locations collected by observers every 30 minutes when following the study groups. We used the minimal and maximal latitude and longitude recorded between 2005 and 2019, producing a rectangular area of 14km*19km. We extracted the mean pixel value for every 8-period in this area, and further compute the monthly means of the daily temperatures (mean pixel values) across 2004-2019.

(iii) Daily rainfall was obtained at 0.25*0.25 degree resolution (corresponding to 28*28km at this latitude) from satellite data sensors (product TRMM 3B42) (Huffman et al. 2016) available on the Giovanni NASA website, using the same geographical area as for temperatures. Monthly accumulated rainfalls (summed across daily values) were subsequently averaged between 2004 and 2019.

(iv) The method of extraction of NDVI, our proxy of food availability, reflecting vegetation cover, is provided in the main text.

Once extracted, we were interested to disentangle seasonal from non-seasonal variations of our four environmental variables, allowing us to test in subsequent analyses the influence of environmental unpredictability on baboon reproduction. These environmental conditions might show either strong seasonality, varying in a consistent predictable pattern across years, or weak seasonality, varying in less predictable way between years. We identified which of these four conditions showed strong versus weak seasonality by assessing how well they were predicted by a sine wave. Sine waves provide an ideal representation of seasonal variation (English et al. 2012; Rickard et al. 2012). At Tsaobis where environmental variation is unimodal, with one rainy season followed by one food peak per year, these variations are by definition periodic (with a period of one year). Importantly, sine waves allow the introduction of a circular variable into a multivariate model: the possible effects of the month or date of birth are circular with a period of one year, and not linear, as the 31$^{st}$ of December is the day before the 1$^{st}$ of January, and should be considered as close as the 31$^{st}$ of October and the 1$^{st}$ of November are, for example. In addition, a sinusoidal term can be used to detect any seasonal effects, i.e. any effect of month or date of birth on reproductive parameters, which may not be captured by our other environmental variables. For example, the phenology of some baboon foods may depend on particular combinations of climatic and photoperiodic cues that very between plant species, and would therefore be difficult to detect using a single environmental predictor. To assess the strength of seasonal variations of each of our environmental factors, we ran four linear models in which each of our four environmental variables was the response variable and a sinusoidal term was the only fixed effect. This sinusoidal term was as follows (as all our environmental variables were monthly values):

$$sin(Month + \varphi)$$

The month of the year in the formula above was converted in a radian measure, so that the period, i.e. one year, equalled to 2*π, ranging from π/6 for January to 2*π for December. We tested different phase values $\varphi$ (0, π/6, π/3, π/2, 2*π/3 or 5*π/6), to account for potential phase shifts across the year. For example, a phase of 0 could maximise the months of March or September, a phase of π/6 could maximise the months of February or August (depending on the sign of the fitted coefficient), etc. We then selected the best phase as the one minimizing the AIC in our model. We found that the sine term of phase π/2, maximising December (the solstice), explains 88% of temperature variation and 99% of photoperiod variation between 2004 and 2019. Thus, in the following analyses, temperature and photoperiod are represented with the sine term only, as we show that between-year variations of temperature and photoperiod are negligible compared to within-year variations. On the contrary, and as expected (see Figure S1, which shows substantial between-year variations of rainfall and NDVI), the sinusoidal term with the best phase only explained 18% of NDVI variation and 20% of rainfall variation. For NDVI and rainfall, we thus decomposed their seasonal and non-seasonal variation. We first computed the mean monthly values across all 15 years, and labelled these variables as 'NDVI_S' and 'Rain_S'. We then computed the difference between each actual monthly value in a given year and this averaged monthly value across years. We labelled these differences as 'NDVI_NS' and 'Rain_NS'. Here, seasonal variables (with the suffix 'S') reflect within-year variation only, i.e., predictable variation, consistent between years; while non-seasonal variables (with the suffix 'NS') reflect between-year variation only, i.e., unpredictable variation, inconsistent between years.

#### Appendix C. Models focusing on interbirth intervals and infant mortality

In these two models, we investigated the costs of environmental variation, group synchrony and rank-related variation of these costs on the fitness of both mothers and offspring, looking at two response variables: interbirth intervals (IBI) and infant mortality before weaning. We tested our three non-exclusive hypotheses, and expected to find fitness costs associated with non-seasonal environmental variations under the non-seasonal environment hypothesis (H1), with reproductive synchrony under the asynchrony hypothesis (H2), and with rank-related fitness costs, possibly linked with environmental fluctuations or synchrony, under the social rank hypothesis (H3). The different fixed effects considered are listed in the main text: non-seasonal environmental variation ('NDVI_NS'), reproductive synchrony ('Number_of_Infants' born around the focal birth), an interaction term between female rank and environmental variation (both seasonal and non-seasonal: 'sin(Date of Birth +$\varphi$)$: Rank'$ and 'NDVI_NS:Rank'), an interaction term between female rank and reproductive synchrony, seasonal environmental variation (sine term of juvenile birth date), the number of adult females in the group ('Number_Adult_Females'), group identity ('Group'), female rank ('Rank'), female parity ('Parity'), and infant sex ('Sex'). We also included female identity ('Identity_female') as a random effect. The two final global models we ran to test our hypothesis are shown in equations (1) and (2) below:

(1) $$IBI \sim NDVI\_NS + Number\_of\_Infants(Window\_IBI) + (\sin(Date\ of\ Birth + \varphi): Rank) + (NDVI\_NS: Rank) + (Number\_of\_Infants(Window\_IBI): Rank) + \sin(Date\ of\ Birth + \varphi) + Number\_Adult\_Females + Group + Rank + Parity + Sex,\ random = Identity\_female,\ \ family = 'gaussian'$$

(2) $Infant\ mortality\ before\ weaning \sim NDVI\_NS + Number\_of\_Infants(Window\_Mortality) + (\sin(Date\ of\ Birth + \varphi){:}\ Rank) + (NDVI\_NS{:}\ Rank) + (Number\_of\_Infants(Window\_IBI){:}\ Rank) + \sin(Date\ of\ Birth + \varphi) + Number\_Adult\_Females + Group + Rank + Parity + Sex,\ random = Identity\_female,\ family = 'binomial'$

In order to run these global models, we first had to identify, among the fixed effects (i) the best sine phase to capture seasonal environmental variation (of six possible phases), (ii) the best measure of non-seasonal environmental variation (of two measures, NDVI_S or rainfall_NS), and (iii) the best time window to capture reproductive synchrony (of nine possible time windows), where the best were defined as those which minimized AIC in the corresponding univariate models. We took as example the IBI models, and we followed the exact same steps for the infant mortality models. Thus:

First, we ran the six following univariate models (3), (4), (5), (6), (7) and (8), considering only the seasonal environmental variations, i.e. the offspring dates of births with a sine term:

(3) $IBI \sim \sin(Date\ of\ Birth), random = Identity\_female, family = 'gaussian'$

(4) $IBI \sim \sin(Date\ of\ Birth + \pi/6), random = Identity\_female, family = 'gaussian'$

(5) $IBI \sim \sin(Date\ of\ Birth + \pi/3), random = Identity\_female, family = 'gaussian'$

(6) $IBI \sim \sin(Date\ of\ Birth + \pi/2), random = Identity\_female, family = 'gaussian'$

(7) $IBI \sim \sin(Date\ of\ Birth + 2 * \pi/3), random = Identity\_female, family = 'gaussian'$

(8) $IBI \sim \sin(Date\ of\ Birth + 5 * \pi/6), random = Identity\_female, family = 'gaussian'$

The only differences between these six models are the value of the phase $\varphi$. For the IBI model, the best phase equaled to π/6 while for infant mortality model, it equaled π/2. By doing so, we characterized the best seasonal environment fluctuations likely to affect our two response variables.

Secondly, we estimated the non-seasonal environmental variation using the NDVI_NS and Rainfall _NS variables described above. We averaged NDVI_NS and Rainfall_NS across the whole period spanning the IBI for Model 1, and from conception to 550 days of age (for live infants) or to death (for dead infants) for Model 2. The two non-seasonal fixed effects: (i) NDVI_NS and (ii) Rain_NS were introduced in separate models given that they reflect the same effect and are well correlated (Pearson correlation test:  R=0.51, t=8.10, p<10-4). We ran the following models, (9) and (10), to determine which non-seasonal environmental variation was the best in each model:

(9) $IBI \sim \sin\left(Date\ of\ Birth + \frac{\pi}{6}\right) + NDVI\_NS, random = Identity\_female, family = 'gaussian'$

(10) $IBI \sim \sin\left(Date\ of\ Birth + \frac{\pi}{6}\right) + Rainfall\_NS, random = Identity\_female, family = 'gaussian'$

For both IBI and infant mortality, the models containing NDVI_NS had lower AIC values. NDVI_S was subsequently used as our metric to represent non-seasonal environmental variation.

Thirdly, we selected the best time window for our reproductive synchrony variable, i.e. the number of infants born around the focal infant (written 'Number_of_Infants' in models (1) and (2)). We used a set of univariate models relying on a strictly similar sample of observations. 'Window_IBI' and 'Window_Mortality' could thus be: before 1 month, after 1 month, both before and after 1 month, before 3 months, after 3 months, both before and after 3 months, before 6 months, after 6 months, or both before and after 6 months. Therefore, we ran nine univariate models for each response variable, with these different time windows, with the structure of the following models (11) and (12):

$$(11) \quad IBI \sim Number\_of\_Infants(Window\_IBI), random = Identity\_female, family = 'gaussian'$$

$$(12) \quad Infant\ mortality\ before\ weaning \sim Number\_of\_Infants(Window\_Mortality), random = Identity\_female, family = 'binomial'$$

We identified the best time window, 'Window_IBI' and 'Window_Mortality': 'Window_IBI' was the number of infants born over the past three months in the same group, while 'Window_Mortality' was the number of infants born six months after in the same group. We finally incorporated these best synchrony time windows and best phase of seasonal variation, along with all other predictors, in our global models (1) and (2).

#### Appendix D. Models focusing on timing of cycle resumption and probability of conception

In these two models, we investigated if females adjusted or delayed their reproductive timings, focusing on their onset of cycle resumption and conception, in order to limit the fitness costs associated with (H1) non-seasonal environmental fluctuations under the non-seasonal environment hypothesis, or (H2) reproductive synchrony under the asynchrony hypothesis. In

(H3), we tested whether female reproductive timings showed rank-related adjustments in response to environmental variations or group synchrony. The different fixed effects considered are listed in the main text: non-seasonal environmental variation (‘NDVI_NS’), reproductive synchrony (‘Mean_Number_of_Conception’ before the reproductive event focal), an interaction term between female rank and environmental variation (both seasonal and non-seasonal: ‘NDVI_S:Rank’ and ‘NDVI_NS:Rank’), an interaction term between female rank and reproductive synchrony (‘Mean_Number_of_Conception:Rank’), seasonal environmental variation (‘NDVI_S’), the number of adult females in the group (‘Number_Adult_Females’), group identity (‘Group’), female rank (‘Rank’) and female parity (‘Parity’). We also included female identity (‘Identity_female’) as a random effect. The two final global models we ran to test our hypothesis are shown in the equations (1) and (2) below:

$$\begin{aligned}
&(1)\; Conception \sim NDVI_NS(Window_NS_Conception) \; + \\
&\quad Mean_Number_of_Conceptions(Window_Conception) + \\
&\quad +(NDVI_S(Window_S_Conception){:}\, Rank) + \\
&\quad (NDVI_NS(Window_NS_Conception) : Rank) + \\
&\quad (Mean_Number_of_Conceptions(Window_Conception){:}\, Rank) + \\
&\quad NDVI_S(Window_S_Conception\,) \; + Number_Adult_Females + \; Group + \\
&\quad Parity + Rank, \; random = Identity_female, family = binomial
\end{aligned}$$

$$\begin{aligned}
&(2)\; Cycle\ resumption \sim NDVI_NS(Window_NS_Cycle_resumption) \; + \\
&\quad Mean_Number_of_Conceptions(Window_Cycle_resumption) + \\
&\quad +(NDVI_S(Window_S_Cycle_resumption){:}\, Rank) + \\
&\quad (NDVI_NS(Window_NS_Cycle_resumption) : Rank) + \\
&\quad (Mean_Number_of_Conceptions(Window_Cycle_resumption){:}\, Rank) + \\
&\quad NDVI_S(Window_S_Cycle_resumption) \; + Number_Adult_Females + \\
&\quad Group + Parity + Rank, \; random = Identity_female, family = binomial
\end{aligned}$$

In order to run these global models, we first had to identify, among the fixed effects (i) the best sine phase to capture seasonal environmental variation (of six possible phases), (ii) the best time window to capture NDVI (NDVI_S and NDVI_NS) and rainfall (Rain_S and Rain_NS) variation, where the best were defined as those which minimized AIC in the corresponding univariate models (iii) the best measure of both seasonal and non-seasonal environmental variation (of three measures of seasonal variations: the sine term, NDVI_S and Rain_S ; and of two measures of non-seasonal variations: NDVI_NS or Rain_NS), and (iv) the best time window to capture reproductive synchrony, where the best were defined as those which minimized AIC in the corresponding multivariate models. The main differences from the steps explained in Appendix 3 are that we considered other aspect of environmental seasonality, and different time windows of group reproductive synchrony. We took as examples the conception models, and we followed the exact same steps for the cycle resumption models.

Before running these models, we first investigated which were the best ecological factors to consider between sinusoidal parameters (seasonal environmental fluctuations only, reflecting temperatures, day time length, or any other seasonal parameter), rainfall and the normalized difference vegetation index (NDVI). The three seasonal effects: (i) the sine wave, (ii) NDVI_S, and (iii) Rain_S were introduced in separate models given that they reflect seasonal environmental variations and were highly correlated (Pearson correlation test: $R>0.84$ and $p<10\text{-}4$ for each pair). Similarly, the two non-seasonal fixed effects: (i) NDVI_NS and (ii) Rain_NS were introduced in separate models given that they reflect the same effect and are well correlated (Pearson correlation test: $R=0.51$, $t=8.10$, $p<10\text{-}4$). We therefore introduced each of our seasonal environmental parameters (sine wave of the month, Rain_S, NDVI_S), and likewise each of our non-seasonal parameters (Rain_NS, NDVI_NS), in separate models. Although we could have simply tested one representative of each, the timing of cycle resumptions and conceptions could be affected by different environmental factors, so we considered the best of all possible environmental predictors for each model.

First of all, we estimated the best phase of the sine fixed effects (with the months in radian), by running the six following models (3), (4), (5), (6), (7), (8):

(3) $Conception \sim \sin(Month)\ ,\ random = Identity\_female, family = binomial$

(4) $Conception \sim \sin(Month + \pi/6)\ ,\ random = Identity\_female, family = binomial$

(5) $Conception \sim \sin(Month + \pi/3)\ ,\ random = Identity\_female, family = binomial$

(6) $Conception \sim \sin(Month + \pi/2)\ ,\ random = Identity\_female, family = binomial$

(7) $Conception \sim \sin(Month + 2 * \pi/3)\ ,\ random = Identity\_female, family = binomial$

(8) $Conception \sim \sin(Month + 5 * \pi/6)\ ,\ random = Identity\_female, family = binomial$

The only differences between these three models are the value of the phase $\varphi$. It was the phase π/6 which was selected. We followed the same method for the timings of cycle resumption as response variables, and we selected π/3 as the best phase.

For rainfall and NDVI fixed effects, we investigated a time window of 0-12 months because (i) other studies found lagged effects of similar length when studying the effect of weather variability on the demography and reproduction of primates (Wiederholt and Post 2011; Campos et al. 2017), and (ii) a lag of more than 12 months (i.e. one annual cycle) would presumably not influence reproductive seasonality. See also Figure S2 for a graphical representation of the 13 time windows tested (taking the example of NDVI_S). We ran sets of univariate models to select the best time window for each rainfall and NDVI predictors (see models (9), (10), (11), (12)). A time window of N months meant that we averaged the value of the fixed effect over the past N months. Therefore, each time window indicated below (‘Rainfall_Window_S_Conception’, ‘Rainfall_Window_NS_Conception’, ‘NDVI_Window_S_Conception’, ‘NDVI_Window_NS_Conception’) reflects the average value of the environmental effect considered over the past N months, N going from 0 to 12.

$$(9)\ Conception \sim Rain\_S(Rainfall\_Window\_S\_Conception),\ random = Identity\_Female, family = binomial$$

$$(10)\quad Conception \sim Rain\_NS(Rainfall\_Window\_NS\_Conception),\ random = Identity\_Female, family = binomial$$

$$(11)\quad Conception \sim NDVI\_S(NDVI\_Window\_S\_Conception),\ random = Identity\_Female, family = binomial$$

$$(12)\quad Conception \sim NDVI\_NS(NDVI\_Window\_NSConception),\ random = Identity\_Female, family = binomial$$

We thus selected Rain_S over the past 4 months, Rain_NS over the past 12 months, NDVI_S over the past 2 months, and NDVI_NS over the past 12 months, respectively. We similarly ran the same models ((9), (10), (11), (12)) with the timings of cycle resumption as the response variable. We selected for the cycle resumption Rain_S over the past 10 months, Rain_NS over the past 4 months, NDVI_S over the past 4 months and NDVI_NS over the past 3 months.

After this first step of identifying the best time window and best phase, we ran other models to estimate which ecological factors, between the sine term, rainfall (Rain_S and Rain_NS), and NDVI (NDVI_S and NDVI_NS) effects, were the best to predict our response variables. To do so, we ran the three following models ((13), (14), and (15)):

$$(13)\quad Conception \sim \sin(Month + pi/3) + (1|ID)$$

$$(14) \quad Conception \sim Rain\_S(4) + Rain\_NS(12) + (1|ID)$$

$$(15) \quad Conception \sim NDVI\_S(2) + NDVI\_NS(12) + (1|ID)$$

We similarly ran models (13), (14) and (15) for cycle resumptions as response variable. For both our response variables, the NDVI model was the best one, and we consequently only kept the NDVI fixed effects in our global model looking at reproductive synchrony effects too (and only presented the models with NDVI fixed effects in the main text of this study).

After selecting the best time windows for ecological factors, and selecting the best ecological factors, we wanted to run a global model considering reproductive synchrony as a fixed effect. The first variable we considered as an indicator of reproductive synchrony was the number of conceptions occurring in the same group. We arbitrarily restricted our exploration to a possible lag of 6 months prior to the focal event, on the basis that females were unlikely to react to reproductive events occurring more than 6 months before. In addition, females in this species were expected to compete over paternal care, which is especially important in the first 6 months of life, the age window in which vulnerability to infanticide is greatest (Palombit 2003), meaning that female reproductive competition may decrease when the age gap between their offspring is greater than 6 months. We therefore investigated the effects of past and present reproductive synchrony by considering (1) a time window of increasing length (from 0 to 6 months) before the observation (model (16) below): here, our fixed effect is the mean number of conceptions in the group occurring in the past X months (X referring to 'Window_Conception' in the model (16) below, and ranging from 0 to 6). See also Figure S2 for a graphical representation of the various past and present time windows for reproductive synchrony that were tested.

$$(16) \quad Conception \sim NDVI\_NS(Window\_NS\_Conception) +$$
$$Mean\_Number\_of\_Conceptions(Window\_Conception) +$$

$$+(NDVI\_S(Window\_S\_Conception){:}\,Rank) +$$

$$(NDVI\_NS(Window\_NS\_Conception) : Rank) +$$

$$(Mean\_Number\_of\_Conceptions(Window\_Conception){:}\,Rank) +$$

$$NDVI\_S(Window\_S\_Conception\ ) \ + Number\_Adult\_Females + \ Group +$$

$$Parity + Rank\ ,\ random = Identity\_Female, family = binomial$$

We therefore ran 7 models for each response variable (see Figure S2). We ran the exact same models, with different best time windows for the NDVI effects, for the cycle resumption response variable. The results shown in the tables of the main text are for those models minimizing AIC, i.e. with the mean number of conceptions over the past four months for the probability of conception, and with the mean number of conceptions over the past six months for the timing of cycling resumption.

The second variable we considered as an indicator of reproductive synchrony was the number of cycling females in the same group. In contrast to the number of conceptions, it was not possible to estimate with precision this number when there were no observers in the field. Therefore, to investigate the potential effects of the number of cycling females in the group (along with our other predictors), we restricted our dataset only to those months where observers were present. Following the same rationale as our other indicator of reproductive synchrony (i.e. the number of conceptions in the group), we investigated the combined effect of current and past synchrony. Nonetheless, due to the limitations of our dataset, we could only explore the effect of reproductive synchrony in the two months prior to a reproductive event. As before, we explored the effect of the mean number of cycling females over the past X months (X referring to 'Window_Cycling_Females' in the model (18), and ranging from 1 to 2), and the effect of the number of cycling females the same month of the observation ('Number_Cycling_Females_Same_Month' in the model (19)).

$$(17)\quad Conception \sim NDVI\_NS(Window\_NS\_Conception) + Mean\_Number\_of\_Cycling\_Females(Window\_Cycling\_Females) + (Mean\_Number\_of\_Cycling\_Females(Window\_Cycling\_Females){:}Rank) + NDVI\_S(Window\_S\_Conception) + Number\_Adult\_Females + Group + Parity + Rank,\ random = Identity\_female, family = binomial$$

$$(18)\quad Conception \sim NDVI\_NS(Window\_NS\_Conception) + Number\_of\_Cycling\_Females\_Same\_Month + (Number\_of\_Cycling\_Females\_Same\_Month{:}Rank) + NDVI\_S(Window\_S\_Conception) + Number\_Adult\_Females + Group + Parity + Rank,\ random = Identity\_female, family = binomial$$

We similarly ran the exact same models, with different best time windows for the NDVI effects, for the cycle resumption response variable.

TABLES

**Table S1**: Different methods used to estimate the dates of 241 conceptions of baboons in Tsaobis between 2005 and 2019.

| Criteria used for estimation | Number of conception estimated | Median of the number of days of uncertainty | Range of the number of days of uncertainty |
|---|---|---|---|
| Conceptions observed in the field | 68 | 0 | 0 |
| Conceptions occurred during a field break | 13 | 10 | 3 - 30 |
| Birth date observed (in the field or during a field break) | 65 | 10 | 10 - 40 |
| Infant coloration & mother’s reproductive state (Dezeure et al. 2021*b*) | 56 | 61 | 0 - 151 |
| Mother’s reproductive state only (with birth observed) | 23 | 65 | 21 - 153 |
| Mother’s reproductive state only (with no birth observed) | 16 | 90 | 24 - 164 |
| Total | 241 | 10 | 0 - 164 |

**Table S2**: Environmental and social determinants on the timing of cycle resumption in a given month, using the number of cycling females as a measure of reproductive synchrony (Model 3.2).

We ran 1000 models with simulated conception date in order to take into account their uncertainty. The table show the mean estimates, mean confidence intervals, mean $X^2$ statistics and mean p-values of the predictors of these 1000 binomial generalized linear mixed model investigating the effect of the number of cycling females in the group on the timing of cycle resumption, along with other fixed effects, based on 61 cycle resumptions from 32 females. Significant effects are indicated in bold. The seasonal environmental variation is the 'NDVI_S' over the past four months. The non-seasonal environmental variation is the 'NDVI_NS' over the past three months. The reproductive synchrony here is characterized by the number of cycling females in the group in the focal month. For categorical predictors, the tested category is indicated between brackets.

| Hypothesis tested | Fixed effects | | Estimate | IC Lower | IC Upper | $X^2$ | P-value |
|---|---|---|---|---|---|---|---|
| H1 | Non-seasonal environmental variation | | 0.25 | 0.01 | 0.49 | 4.50 | 0.064 |
| H2 | Reproductive synchrony | | 0.05 | -0.23 | 0.32 | 0.29 | 0.685 |
| Control | **Seasonal environmental variation** | | **-0.30** | **0.56** | **-0.05** | **5.49** | **0.031** |
| | Number of adult females | | -0.10 | -0.45 | 0.25 | 0.41 | 0.576 |
| | Group | (L) | 0.11 | -0.44 | 0.66 | 0.59 | 0.763 |
| | | (M) | -6.71 | -949.95 | 936.52 | | |
| | Rank | | -0.09 | -0.35 | 0.17 | 0.58 | 0.517 |
| | Parity | (primiparous) | -0.12 | -0.82 | 0.58 | 0.25 | 0.693 |

**Table S3:** Environmental and social determinants on the probability of conception in a given month, using the number of conceptions as a measure of reproductive synchrony (Model 4.2).

We ran 1000 models with simulated conception date in order to take into account their uncertainty. The table show the mean estimates, mean confidence intervals, mean $X^2$ statistics and mean p-values of the predictors of these 1000 binomial generalized linear mixed model investigating the effect of the number of cycling females in the group on the probability of conception, along with other fixed effects, based on 103 conceptions out of 759 observations from 50 females. Significant effects are indicated in bold. For relevant significant effect, we also indicated in the footnote the Wilcoxon confidence interval of the p-values. The seasonal environmental variation is the 'NDVI_S' over the past two months. The non-seasonal environmental variation is the 'NDVI_NS' over the past 12 months. The reproductive synchrony here is characterized by the number of cycling females in the group in the focal month. For categorical predictors, the tested category is indicated between brackets.

| Hypothesis tested | Fixed effects | | Estimate | IC | | $X^2$ | P-value |
|---|---|---|---|---|---|---|---|
| | | | | Lower | Upper | | |
| H1 | Non-seasonal environmental variation | | -0.21 | -0.43 | 0.01 | 3.64 | 0.128 |
| H2 | **Reproductive synchrony** | | **-0.25** | **-0.48** | **-0.01** | **4.43** | **0.050**[a] |
| Control | Seasonal environmental variation | | 0.17 | -0.04 | 0.37 | 2.72 | 0.128 |
| | Number of adult females | | 0.12 | -0.23 | 0.47 | 0.50 | 0.514 |
| | Group | (L) | -0.15 | -0.63 | 0.34 | 0.97 | 0.654 |
| | | (M) | 0.09 | -62.8 | 63.97 | | |
| | Rank | | 0.04 | -0.19 | 0.27 | 0.19 | 0.720 |
| | Parity | (nulliparous) | -0.64 | -1.17 | -0.11 | 6.24 | 0.053 |
| | | (primiparous) | 0.50 | -0.62 | 0.72 | | |

[a] CI: [0.0413 – 0.0460]

**Table S4**: Effect of the number of conceptions over various time-windows on the probability to conceive on a given month.

For each modality of reproductive synchrony time window (mean number of conception over the past 0-6 months), we ran (i) 1000 models without any rank interactions, and extracted the mean AIC of these models, along with estimate and P-value of the reproductive synchrony fixed effect ('Term alone' rows) ; and (ii) 1000 full models with rank interactions (see Table 5), and extracted the mean AIC of these models, along with estimate and P-value of the interaction between female rank and reproductive synchrony fixed effect ('Interaction rank' rows). The best model fits, with and without the rank interaction, are shown in bold writing.

| Group reproductive synchrony | AIC | Fixed effects | Estimates | P-values |
|---|---|---|---|---|
| Number of conception the same month | 1145.1 | Term alone | 0.09 | 0.270 |
| | 1148.1 | Interaction Rank | -0.01 | 0.160 |
| Mean number of conception over last month | 1142.6 | Term alone | 0.16 | 0.053 |
| | 1143.6 | Interaction Rank | -0.16 | 0.043 |
| Mean number of conception over the last 2 months | 1140.1 | Term alone | 0.20 | 0.010 |
| | 1142.1 | Interaction Rank | -0.14 | 0.059 |
| Mean number of conception over the last 3 months | 1140.6 | Term alone | 0.21 | 0.013 |
| | 1142.3 | Interaction Rank | -0.16 | 0.046 |
| **Mean number of conception over the last 4 months** | **1136.7** | **Term alone** | **0.26** | **0.002** |
| | **1137.0** | **Interaction Rank** | **-0.18** | **0.021** |
| Mean number of conception over the last 5 months | 1138.5 | Term alone | 0.24 | 0.004 |
| | 1139.9 | Interaction Rank | -0.17 | 0.036 |
| Mean number of conception over the last 6 months | 1138.9 | Term alone | 0.25 | 0.005 |
| | 1141.0 | Interaction Rank | -0.16 | 0.057 |

**Table S5**: Effect of the number of conceptions over various time-windows on the timing of cycle resumption on a given month.

For each modality of reproductive synchrony time window (mean number of conception over the past 0-6 months), we ran (i) 1000 models without any rank interactions, and extracted the mean AIC of these models, along with estimate and P-value of the reproductive synchrony fixed effect ('Term alone' rows) ; and (ii) 1000 full models with rank interactions (see Table 4), and extracted the mean AIC of these models, along with estimate and P-value of the interaction between female rank and reproductive synchrony fixed effect ('Interaction rank' rows). The best model fits, with and without the rank interactions, are shown in bold writing.

| Group reproductive synchrony | AIC | Fixed effects | Estimates | P-values |
|---|---|---|---|---|
| Number of conception the same month | 1008.3 | Term alone | 0.01 | 0.597 |
| | 1011.8 | Interaction Rank | 0.07 | 0.473 |
| Mean number of conception over last month | 1008.5 | Term alone | -0.06 | 0.529 |
| | 1011.9 | Interaction Rank | 0.09 | 0.402 |
| Mean number of conception over the last 2 months | 1007.0 | Term alone | -0.15 | 0.177 |
| | 1010.1 | Interaction Rank | 0.10 | 0.332 |
| **Mean number of conception over the last 3 months** | **1005.7** | **Term alone** | **-0.19** | **0.084** |
| | 1008.7 | Interaction Rank | 0.12 | 0.275 |
| Mean number of conception over the last 4 months | 1007.9 | Term alone | -0.13 | 0.236 |
| | 1010.4 | Interaction Rank | 0.13 | 0.204 |
| Mean number of conception over the last 5 months | 1008.3 | Term alone | -0.12 | 0.253 |
| | 1009.7 | Interaction Rank | 0.17 | 0.109 |
| **Mean number of conception over the last 6 months** | 1007.2 | Term alone | -0.17 | 0.123 |
| | **1006.8** | **Interaction Rank** | **0.22** | **0.035** |

### FIGURES

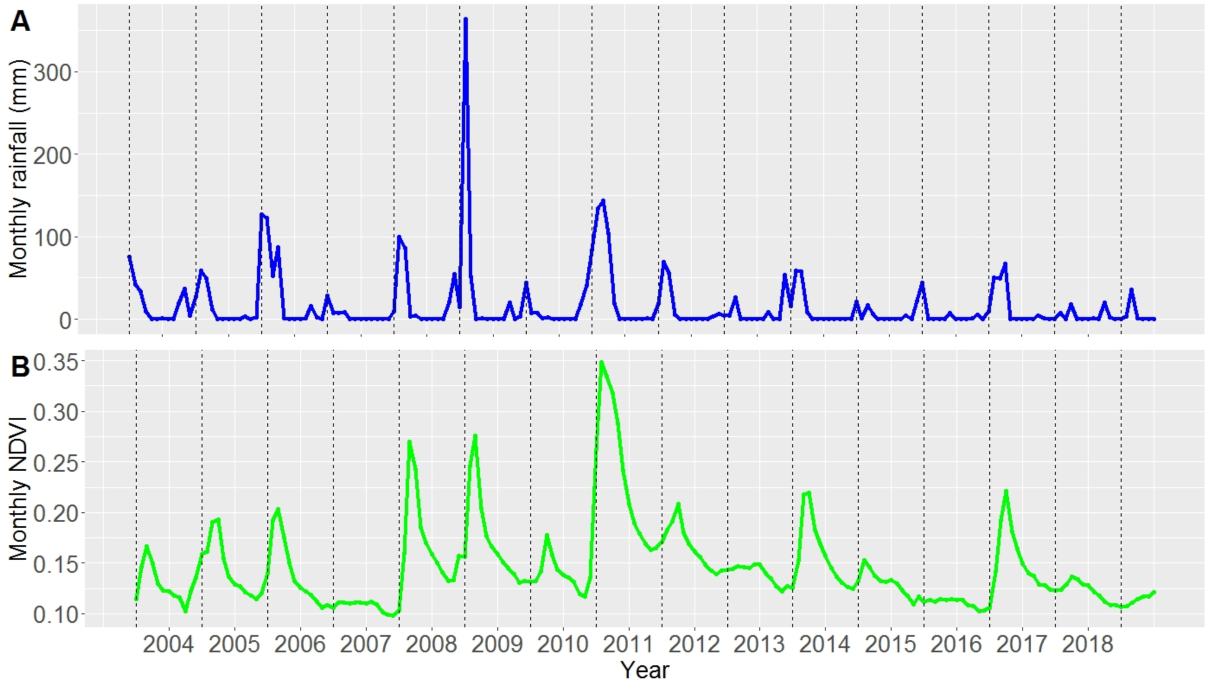

**Figure S1**: Tsaobis rainfall and food availability variations over the study period (2004-2019).

We represented the monthly cumulative rainfall in mm (Panel A) and monthly NDVI value on J group homerange (Panel B) according to time (between January 2004 and July 2019). Rainfall and NDVI values were extracted from satellite data, resp. from the Giovanni NASA website (TRMM 3B42 product) (Huffman et al. 2016) and the MODIS NASA dataset (MODIS13A1 product) (see also Appendix B and Dezeure et al. 2021*a*). We indicated with vertical dashed black line the month of January for each year.

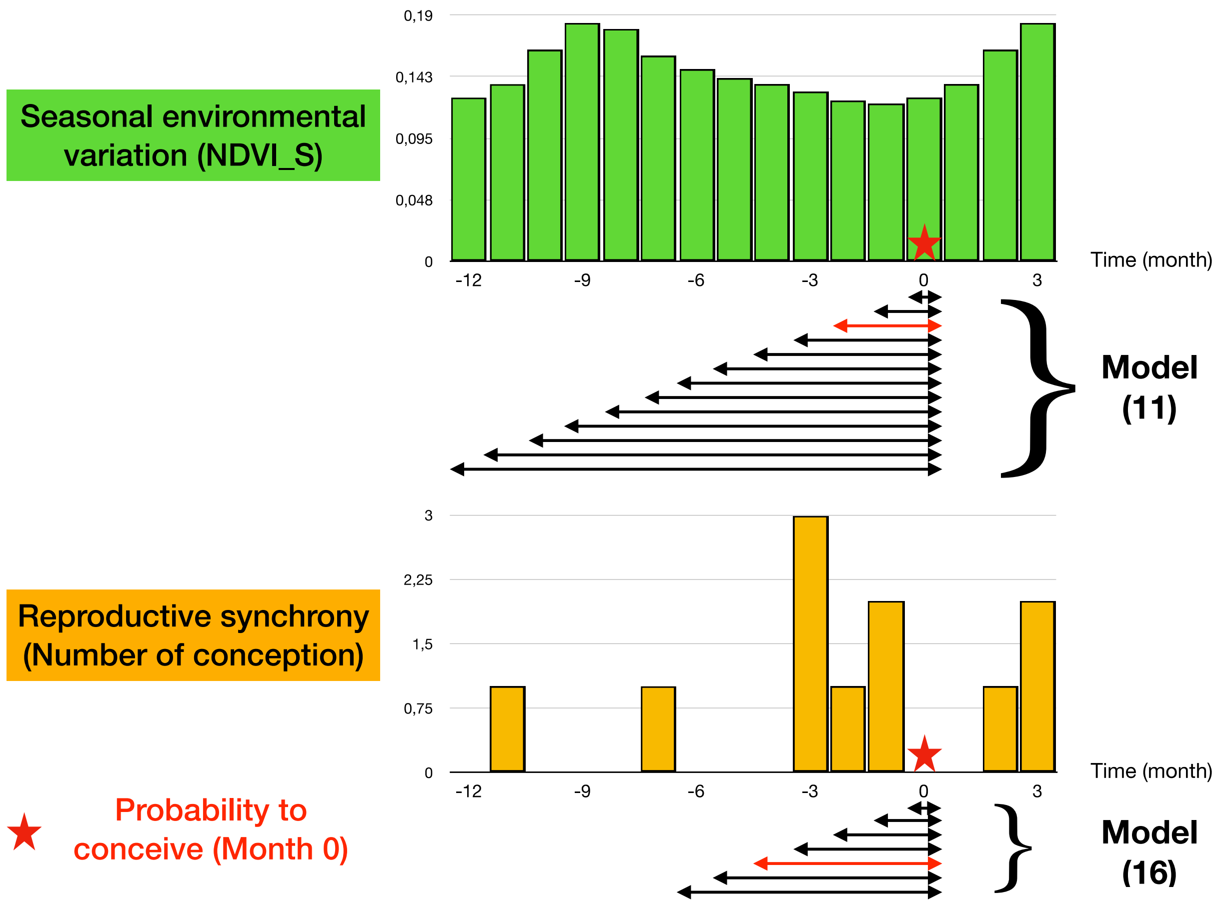

**Figure S2**: Examples of the different time windows considered when investigating the effects of current and past environmental variation and reproductive synchrony on female conception probability.

The model numbers indicated correspond to the ones listed in Appendix B. We plotted the seasonal NDVI ('NDVI_S') per month in green and the number of conceptions in J group per month in yellow, considering December 2018 as time 0 (and so, for example, December 2017 is indicated with -12). Model (11) tested the effect of the mean NDVI_S over the X past months, X going from 0 to 12, and was thus run 13 times. Model (16) tested the effect of the mean number of conceptions over the past X months, X going from 0 to 6, and was thus run seven times. The arrows in red indicate the best time window effect selected for each model sets, with conception probability as response variable.

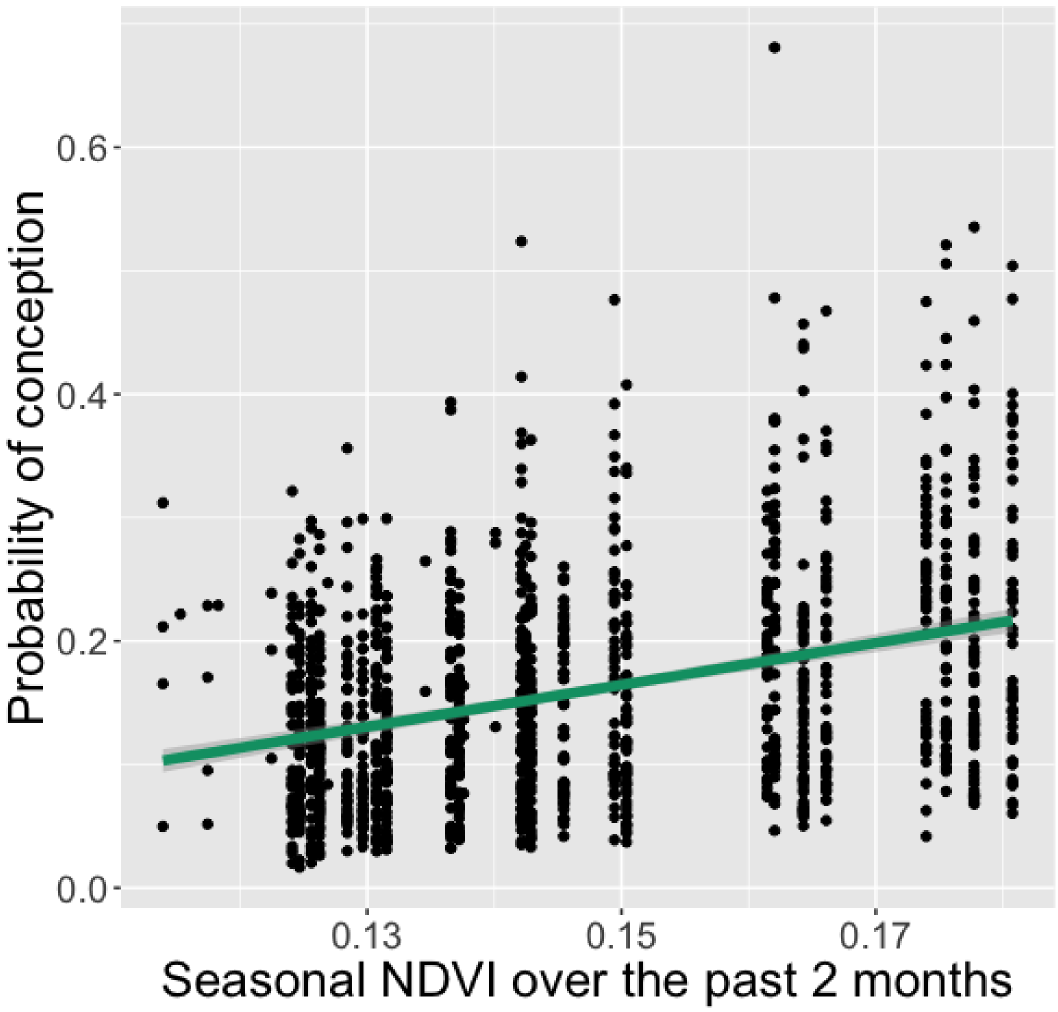

**Figure S3**: The probability of conception increases with higher seasonal variation of NDVI over the past two months.

Each black dots represents a fitted value of the full model (Table 5) focusing on the probability of conception according to the mean seasonal NDVI over the past 2 months. The green curve shows the logistic fit (using the glm method of stat_smooth function), and the shaded area displays 95% confidence intervals around it.
